# Bone Material Properties Distributions from quantitative Raman analysis: a distribution-based approach to human bone matrix characterization

**DOI:** 10.64898/2026.09.10.750589

**Authors:** Nadine Gaborit, Stephanie Lemière, Aleksandra Mieczkowska, Guillaume Mabilleau

**Author notes:** Contributed equally to this work (Nadine Gaborit, Stephanie Lemière, Aleksandra Mieczkowska). Corresponding author: Guillaume Mabilleau, Inserm UMR_S 1229 RMeS – REGOS, Université d’Angers, Institut de Biologie en Santé, 4 rue Larrey, F-49933 Angers, France.

## Abstract

**Background:** Bone mechanical competence depends not only on bone mass but also on the spatial organization of mineral and organic matrix properties. Whereas quantitative backscattered electron imaging (qBEI) characterizes mineral heterogeneity using Bone Mineral Density Distributions (BMDDs), no equivalent framework currently exists for describing the biochemical heterogeneity of the bone matrix by Raman microspectroscopy.

**Methods:** We developed a quantitative Raman analysis (qRA) workflow to generate Bone Material Properties Distributions (BMPDs). Raman spectra were acquired along full-thickness line scans crossing successive bone structural units in human trabecular and cortical bone. Pixel-wise Raman-derived biochemical parameters describing collagen organization, advanced glycation end-products, glycosaminoglycans, mineral-to-matrix ratios, carbonate substitution, and mineral crystallinity were converted into normalized frequency distributions and modeled using gaussian functions. BMPDs were generated from undecalcified pMMA-embedded iliac biopsies (n=38) and routinely processed decalcified paraffin-embedded osteomedullary biopsies (n=21). The analytical framework was validated against conventional two-dimensional Raman mapping, and reproducibility, interindividual variability, and demographic influences were evaluated.

**Results:** BMPDs were consistently well described by gaussian functions (typically R^2^ ≥ 0.95), allowing extraction of three descriptors for each biochemical parameters: Mean, Peak, and Width. One-dimensional line scans showed excellent agreement with conventional two-dimensional Raman mapping while reducing acquisition time approximately 100-fold. Technical variability remained substantially lower than biological variability for all parameters, and a single trabecular line scan provided reliable estimates of BMPD descriptors for nearly all Raman-derived indices. Cortical bone exhibited narrower BMPDs than trabecular bone, indicating lower biochemical heterogeneity. Comparable BMPDs were obtained from undecalcified and routinely decalcified specimens, demonstrating robustness to tissue processing. No significant associations with age or sex were observed after correction for multiple testing. Reference BMPDs further enabled visualization and standardized quantification of deviations in representative pathological bone biopsies.

**Conclusions:** Quantitative Raman analysis introduces BMPDs as a novel framework for assessing the spatial heterogeneity of bone extracellular matrix composition. By extending Raman microspectroscopy beyond conventional mean measurements, BMPDs provide robust, reproducible descriptors of bone material organization and establish a methodological foundation for investigating alterations of bone quality in metabolic bone diseases.

## 1. Introduction

Bone strength depends on the coordinated organization of bone quantity, microarchitecture and tissue material properties. At the tissue level, the extracellular matrix is a composite material consisting of a mineral phase embedded within an organic matrix primarily composed of type I collagen and non-collagenous proteins ^(1)^. Alterations in either component influence bone mechanical competence and contribute to skeletal fragility independently of bone mass.

Over the past three decades, quantitative backscattered electron imaging (qBEI) has become one of the reference techniques for characterizing bone mineralization at the tissue level ^(2)^. Rather than relying solely on mean calcium content, qBEI introduced the concept of Bone Mineral Density Distribution (BMDD), in which the frequency distribution of mineralization values provides quantitative descriptors of both mineral content and tissue heterogeneity. This distribution-based approach has substantially improved our understanding of bone remodeling, mineralization kinetics and bone diseases.

In contrast, no equivalent analytical framework currently exists for characterizing the biochemical heterogeneity of the organic bone matrix. Raman microspectroscopy has emerged as a powerful technique for assessing bone composition because it simultaneously provides information on mineral and collagen components at micrometer spatial resolution without requiring exogenous labeling. Numerous Raman-derived parameters have been proposed to evaluate collagen secondary structure, collagen maturity, post-translational modifications and extracellular matrix composition ^(3)^. However, these parameters are generally summarized as mean values over a region of interest, often between double calcein or tetracycline labels or at the bone periosteal surface ^(4–6)^, thereby overlooking the spatial heterogeneity generated by bone remodeling and the succession of bone structural units (BSUs). Because each BSU corresponds to a distinct remodeling event and tissue age, the biochemical properties of the organic matrix vary continuously across bone tissue. Consequently, the distribution of Raman-derived parameters may contain biologically relevant information that is not captured by conventional averaging approaches. Inspired by the BMDD concept, we hypothesized that describing the statistical distribution of Raman-derived biochemical parameters would provide robust descriptors of bone matrix organization and heterogeneity.

In the present study, we introduce a quantitative Raman analysis (qRA) workflow for the generation of Bone Material Properties Distributions (BMPDs). The method consists of acquiring Raman spectra along linear profiles crossing all BSUs within individual trabeculae or cortical, calculating biochemical Raman parameters at each micrometer, and modeling their distributions using Gaussian fitting. Similar to BMDD analysis, the resulting distributions are characterized by their mean value, peak position and full width at half maximum, providing quantitative descriptors of both matrix composition and heterogeneity. As a proof of concept, the method was evaluated using osteomedullary and transiliac bone biopsies obtained from individuals without skeletal fragility. To assess its robustness across commonly used histological preparations, BMPDs were generated from both polymethyl methacrylate (pMMA)-embedded undecalcified bone and decalcified paraffin-embedded bone sections. This study establishes the methodological framework of qRA-derived BMPDs and provides the basis for future investigations of bone matrix alterations in metabolic bone diseases.

## 2. Materials and methods

### 2.1. Patients and bone biopsy preparation

The methodological validation of the quantitative Raman analysis (qRA) workflow was performed using two independent sets of human bone specimens prepared according to standard histopathological and histomorphometric procedures. Undecalcified transiliac bone biopsies obtained at necropsy from individuals without evidence of skeletal fragility or metabolic bone disease, previously reported by Zhioua et al. ^(7)^, were used for the analysis of mineralized bone. This cohort consisted of 22 men and 16 women aged 20 to 74 years (median age, 51 years; 95% CI, 33–61 years). These biopsies were obtained using an 8-mm inner diameter trephine, as commonly performed for the diagnosis of metabolic bone disease. Following fixation in 80% ethanol, specimens were dehydrated in graded ethanol series and embedded in polymethylmethacrylate (pMMA) without prior decalcification. PMMA blocks were subsequently ground using SiC paper (320–4000 grit) and polished with diamond paste (1 µm) (Struers, Champigny-sur-Marne, France). Raman analyses were performed directly at the surface of the pMMA block. For the decalcified cohort, commercially available histological sections (5µm-thick) derived from osteomedullary biopsies were used. These samples originated from the BM481a tissue microarray (TMA, Lot# 210917, Biomax.us now TissueArray.com LLC, Derwood, MD) and were distributed as fully processed paraffin-embedded sections by Euromedex (Souffelweyersheim, France). Among the 24 individuals represented in the TMA, 21 contained at least one trabecular bone core in one of the duplicate suitable for Raman acquisition and were included in the analysis. This cohort consisted of 18 men and 3 women aged 19 to 72 years (median age, 60 years; 95% CI, 50–64 years). The corresponding biopsies had been routinely fixed in neutral phosphate-buffered formalin within 30 minutes of surgical excision, decalcified with nitric acid and embedded in paraffin prior to tissue microarray construction, following standardized protocols used in clinical pathology laboratories ^(8,9)^. The resulting sections (1.5-mm cores in duplicate) were mounted on mirror-polished stainless-steel slides specifically optimized for Raman microspectroscopy, in accordance with previously published recommendations ^(10)^. In addition, a transiliac biopsy from a 58 year-old male with osteomalacia secondary to phosphate diabetes in a renal transplant recipient (see Sayegh et al. ^(11)^ for more information on this patient) and an osteomedullary biopsy from a 70 year-old male with a multiple myeloma (see Josselin et al.^(12)^ for patient details) were also evaluated.

### 2.2. Raman acquisition (qRA)

Quantitative Raman analysis (qRA) was performed using an inVia Qontor confocal Raman microspectrometer (Renishaw Plc, Wotton-under-Edge, UK) equipped with a 785-nm diode laser, a 20× objective (numerical aperture 0.40), and a 1200 lines/mm diffraction grating, providing a spectral resolution of approximately 1 cm^−1^. The laser power at the sample was 30 mW, and the laser spot diameter was approximately 2.4 µm. At the beginning of each acquisition day, the Raman microspectrometer was calibrated by performing spectral calibration using the internal silicon standard and laser beam alignment. For each trabecula, Raman spectra were acquired along a linear profile positioned perpendicular to the longitudinal axis of the trabecula, extending from one trabecular surface to the opposite surface. This acquisition strategy was designed to intersect all successive bone structural units (BSUs) present within the trabecula and to capture the spatial heterogeneity of the bone extracellular matrix. For undecalcified pMMA-embedded specimens, an additional linear acquisition was performed on cortical bone regions. In these cases, the line scan was positioned across the cortical thickness to encompass multiple osteons and interstitial bone tissue, thereby intersecting several cortical BSUs within a single acquisition profile. Spectra were collected every micrometer using point acquisitions with an integration time of 5 s and three accumulations per measurement, over the spectral range of 800–1800 cm^−1^. For paraffin-embedded and pMMA-embedded specimens, reference spectra of pure paraffin and pure pMMA, respectively, were acquired under identical instrumental conditions. These reference spectra were subsequently used during spectral preprocessing to remove the spectral contribution of the corresponding embedding medium by spectral subtraction.

### 2.3. Spectral preprocessing

All Raman spectra were processed using a standardized preprocessing pipeline implemented in custom scripts developed in MATLAB R2025b (MathWorks, Natick, MA, USA). First, fluorescence background was corrected by fitting and subtracting a fifth-order polynomial baseline from each spectrum. Baseline-corrected spectra were subsequently normalized using the standard normal variate (SNV) transformation to reduce multiplicative intensity variations arising from acquisition conditions and sample-to-sample variability. Spectral noise was then reduced using a Savitzky-Golay filter (second-order polynomial, 17-point window), preserving the shape and intensity of the Raman bands. Finally, the residual contribution of the embedding medium was then removed using a weighted Extended Multiplicative Signal Correction (EMSC) approach. The EMSC model included constant, linear and quadratic baseline terms together with a reference spectrum of pure paraffin (or pure pMMA) and its first derivative to account for slight spectral shifts. Model fitting was performed using weighted least squares, with the CH_2_ region (1400-1470 cm^−1)^ excluded from the optimization to avoid overcorrection of bone-derived Raman bands.

Following preprocessing, Raman-derived biochemical parameters were calculated for each bone pixel using peak intensity ratios. Organic matrix parameters were determined in both paraffin-embedded decalcified specimens and undecalcified pMMA-embedded specimens and included:

- the 1670/1640 cm^−1^ ratio, reflecting collagen secondary structure ^(13)^;
- the 1670/1690 cm^−1^ ratio, reflecting collagen maturity ^(13)^;
- the Hyp/Pro (~872/~854 cm^−1)^ ratio, reflecting proline hydroxylation ^(14)^;
- the CML/CH_2_ (~1150/~1450 cm^−1)^ ratio, reflecting carboxymethyllysine content ^(15)^;
- the PEN/CH_2_ (~1495/~1450 cm^−1)^ ratio, reflecting pentosidine content ^(15)^;
- the GAG/Amide III (~1378/~1250 cm^−1)^ ratio, reflecting glycosaminoglycan content relative to the collagen matrix ^(16,17)^.

For undecalcified PMMA-embedded specimens only, additional parameters describing the mineral phase were calculated, including:

- ν_1_PO_4_/Amide I (~960/~1650 cm^−1)^, reflecting the mineral-to-collagen ratio ^(5)^;
- ν_1_PO_4_/Amide III (~960/~1250 cm^−1)^, reflecting the mineral-to-collagen ratio ^(5)^;
- ν_1_PO_4_/CH_2_ (~960/~1450 cm^−1)^, reflecting the mineral-to-organic matrix ratio ^(5)^;
- ν_1_PO_4_/Hyp–Pro (~960/(~872+~854) cm^−1)^, reflecting the mineral-to-collagen ratio ^(5)^;
- CO_3_/PO_4_ (~1070/~960 cm^−1)^, reflecting carbonate substitution within the apatite crystal lattice ^(18)^;
- mineral crystallinity, calculated as the inverse of the full width at half maximum of the ν_1_PO_4_ band (~960 cm^−1)^, reflecting apatite crystal size and structural order ^(19,20)^.

For pMMA-embedded cortical bone, Haversian canals and osteocyte lacunae were excluded prior to BMPD construction. These non-mineralized regions were identified using the 812 cm^−1^/ν_1_PO_4_ intensity ratio calculated before pMMA spectral subtraction. Pixels with an 812 cm^−1^/ν_1_PO_4_ ratio≥1, indicating predominance of embedding medium over mineralized tissue, were classified as non-bone and excluded from subsequent analyses. In trabecular bone, acquisition lines were manually positioned to avoid visible osteocyte lacunae whenever possible. The resulting pixel-wise Raman values constituted the input dataset for BMPD construction, with one distribution generated for each Raman-derived parameter.

### 2.4. Histogram distribution and BMPD construction

For each Raman-derived biochemical parameter, the pixel-wise values obtained along an individual trabecular or cortical line scan were used to construct a BMPD. For each specimen, pixel values were grouped into 100 equally spaced bins spanning the full range of observed values for the corresponding parameter. The use of 100 bins provided sufficient resolution to describe the distribution while maintaining stable estimates of the relative frequency in each interval and is analogous to the construction of BMDD in qBEI ^(2)^. The lower and upper boundaries of the binning range were empirically determined through preliminary testing to ensure an adequate spread of pixel values across the histogram, avoiding excessive clustering of pixels within a single bin while preserving the overall distribution shape. The number of pixels within each bin (n_i_) was expressed as a percentage of the total number of bone pixels sampled along the acquisition line, yielding a normalized frequency distribution (∑n_i_ =100%). The resulting normalized histogram constituted the BMPD of the investigated Raman parameter and described its spatial distribution across all bone pixels intersected by the line scan.

### 2.5. Gaussian modeling and extracted parameters

Each BMPD was modeled using a Gaussian function according to Equation (1), where A represents the maximum amplitude, μ the position of the distribution peak, and σ the standard deviation

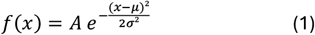

The gaussian fit was used as a mathematical model to summarize the overall shape of each BMPD, following the conceptual framework established for BMDD analysis. Goodness of fit was quantified using the coefficient of determination (R^2^), which was used to evaluate how closely each BMPD approximated a gaussian distribution. Because the quality of the fit may depend on both the intrinsic distribution of the biochemical parameter and the number of Raman measurements contributing to the histogram, R^2^ values were analyzed descriptively and were not used as an exclusion criterion for subsequent analyses. As in qBEI, the fitted gaussian was used solely to derive descriptors of the distribution. Only μ and σ were retained for biological interpretation, whereas the gaussian amplitude A was not considered an independent tissue descriptor. For each BMPD, three primary descriptors were derived from the gaussian fit:

- *BMPD_peak*: corresponded to the position of the distribution peak, µ, representing the most prevalent value of the Raman-derived parameter within the bone segment.
- *BMPD_width*: defined as the full width at half maximum of the fitted distribution and computed according to Equation (2). Width was used as an indicator of spatial heterogeneity of the corresponding bone material property across the analyzed bone structural units.

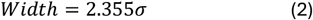
- *BMPD_mean*: corresponded to the weighted arithmetic mean of pixel-wise Raman values within the acquisition line describing the average biochemical composition, calculated from histogram data according to Equation (3) where x_i_ is the bin center and f_i_ the percentage of pixels within bin *i*.

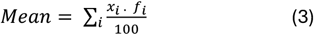

### 2.6. Comparison of two-dimensional ROI and one-dimensional line-scan Raman sampling

To evaluate whether one-dimensional line-scan Raman acquisition provides equivalent information to conventional two-dimensional sampling, a spatial validation analysis was performed. Square regions of interest (ROIs) (100 µm × 100 µm), encompassing the full trabecular width, were defined on trabecular bone sections, and full pixel-wise Raman maps were acquired within each ROI. Pixels with an 812/PO_4_ ratio≥1 were classified as non-bone and excluded from subsequent analyses. This design ensured that the entire trabecular thickness and associated bone structural units were captured within a single field of view. A centered linear profile was then extracted from each ROI, passing through its geometric center. This line scan was used to simulate reduced-dimensional sampling while preserving identical spatial localization. The analysis was performed on 12 randomly selected pMMA-embedded undecalcified biopsies and the 21 paraffin-embedded decalcified biopsies, ensuring balanced representation of both tissue preparation conditions. BMPD_mean, BMPD_peak, and BMPD_width were calculated independently from ROI-based and line-scan–based datasets. Both approaches were applied to the same anatomical regions in order to ensure direct comparability of spatial sampling strategies.

### 2.7. Evaluation of the impact of decalcification and tissue processing on BMPD measurements

To assess whether the decalcification and tissue processing workflow affected Raman-derived biochemical parameters, measurements obtained from decalcified paraffin-embedded bone specimens were compared with those obtained from the undecalcified pMMA-embedded bone biopsies. Undecalcified bone biopsies embedded in pMMA were sectioned at a thickness of 7 µm using a microtome. Sections were subsequently flattened and transferred onto stainless steel slides to enable decalcification while preserving tissue morphology and spatial organization. Decalcification was performed using 10% ethylenediaminetetraacetic acid (EDTA) overnight at 4°C. Following decalcification, Raman measurements were acquired on trabecular bone regions using the previously established 1D line-scan acquisition approach. Raman-derived biochemical parameters were calculated and compared with those obtained from the paraffin reference cohort to evaluate the impact of the decalcification and processing workflow on BMPD measurements.

### 2.8. Statistical analysis

Statistical analyses were performed using GraphPad Prism 11 (GraphPad Software, Boston, MA, USA), unless otherwise specified. Continuous variables are reported as mean ± standard deviation (SD). Agreement between one-dimensional Raman line-scan and two-dimensional region-of-interest acquisitions was assessed using three complementary approaches. The monotonic relationship between both acquisition strategies was evaluated using Spearman’s rank correlation coefficient (ρ), whereas method agreement and potential systematic bias were assessed using Bland– Altman analysis. The reliability of BMPD descriptors obtained from both acquisition approaches was further quantified using the intraclass correlation coefficient (ICC), calculated using a two-way mixed-effects model for absolute agreement. ICC analyses were performed in R software (version 4.6.1; R Foundation for Statistical Computing, Vienna, Austria), while all other statistical analyses were conducted using GraphPad Prism. Statistical significance was defined as p < 0.05. To determine the minimum number of Raman line scans required to obtain a reproducible estimate of BMPD descriptors, variance component analyses were performed using three independent full-thickness Raman line scans acquired across distinct trabecular or cortical regions from 15 biopsies. For each BMPD descriptor, a linear mixed-effects model was fitted using the lme4 package in R, with patient included as a random effect. The model partitioned the total variance into between-patient (τ^2^) and within-patient (σ^2^) components. The single measure intraclass correlation coefficient (ICC(1)) and its 95% confidence interval were derived from these variance components using 500 parametric bootstrap simulations. The Spearman– Brown prophecy formula was subsequently applied to estimate the number of independent Raman line scans required to achieve a predefined reliability threshold of ICC ≥ 0.80. To investigate the potential influence of demographic characteristics on BMPD descriptors, linear regression models were fitted independently for each Raman-derived biochemical parameter and descriptor using R software. Age and sex were included as explanatory variables, and BMPD descriptors were used as dependent variables. Analyses were performed separately for each tissue preparation group: decalcified paraffin-embedded trabecular bone, undecalcified pMMA-embedded trabecular bone, and undecalcified pMMA-embedded cortical bone. Associations were evaluated using regression coefficients (β), R^2^, and adjusted R^2^ values. P-values were corrected for multiple comparisons using the Benjamini–Hochberg false discovery rate (FDR) procedure, and an adjusted p-value < 0.05 was considered statistically significant. For exploratory assessment of pathological bone samples, individual BMPDs were compared descriptively with the corresponding reference BMPD distributions obtained from the control cohorts. Deviations from reference distributions were quantified using standardized differences (Z-scores) calculated for BMPD descriptors. Because pathological analyses were based on single-case observations, no formal statistical hypothesis testing was performed. To determine whether Raman spectral descriptors obtained from purchased decalcified paraffin-embedded osteomedullary biopsies were equivalent to those obtained from EDTA-decalcified, pMMA-embedded iliac bone biopsies, a Shapiro-Wilk test was performed to assess normality and when passed an unpaired t-test was performed. When normality was not respected, the Mann-Whitney test was used. Statistical significance was defined as p < 0.05.

## 3. Results

### 3.1. Construction of BMPDs and Gaussian modeling

BMPDs were developed as the Raman microspectroscopy equivalent of BMDD, providing a quantitative framework to describe the spatial distribution of bone material properties. The workflow used to generate BMPD is illustrated in Figure 1. Briefly, Raman-derived biochemical parameter values obtained from all bone pixels along each line scan were pooled to generate normalized frequency distributions for each bone material parameter. This approach was applied to both decalcified paraffin-embedded bone sections and undecalcified pMMA-embedded polished specimens. Figure 2 illustrates the construction of a representative BMPD for the 1670/1690 ratio from a trabecula in a paraffin-embedded bone biopsy. Raman spectra acquired along a randomly positioned line scan were used to calculate the pixel-wise 1670/1690 ratio, from which the corresponding BMPD was generated. Gaussian fitting accurately described the experimental distribution (R^2^ = 0.961), yielding a trabecular 1670/1690_mean of 1.41, a 1670/1690_peak of 1.40, and a 1670/1690_width of 0.220. Comparable BMPDs were obtained for additional Raman-derived parameters reflecting collagen secondary structure, proline hydroxylation, glycosaminoglycan content, carboxymethyllysine content, and pentosidine content (Supplementary Figure S1). The same analytical workflow was applied to undecalcified pMMA-embedded cortical and trabecular bone (Figure 3). In cortical bone, the 1670/1690 BMPD was well described by a Gaussian model (R^2^ = 0.998), with 1670/1690_mean, 1670/1690_peak, and 1670/1690_width values of 1.52, 1.52, and 0.229, respectively. In trabecular bone, the corresponding values were 1.48, 1.48, and 0.187, with an R^2^ of 0.997. BMPDs generated for additional organic and mineral matrix parameters also exhibited excellent agreement with gaussian fits in both cortical and trabecular bone (Supplementary Figures S2 and S3), with most fits yielding R^2^ values ≥ 0.95.

**Figure 1.**
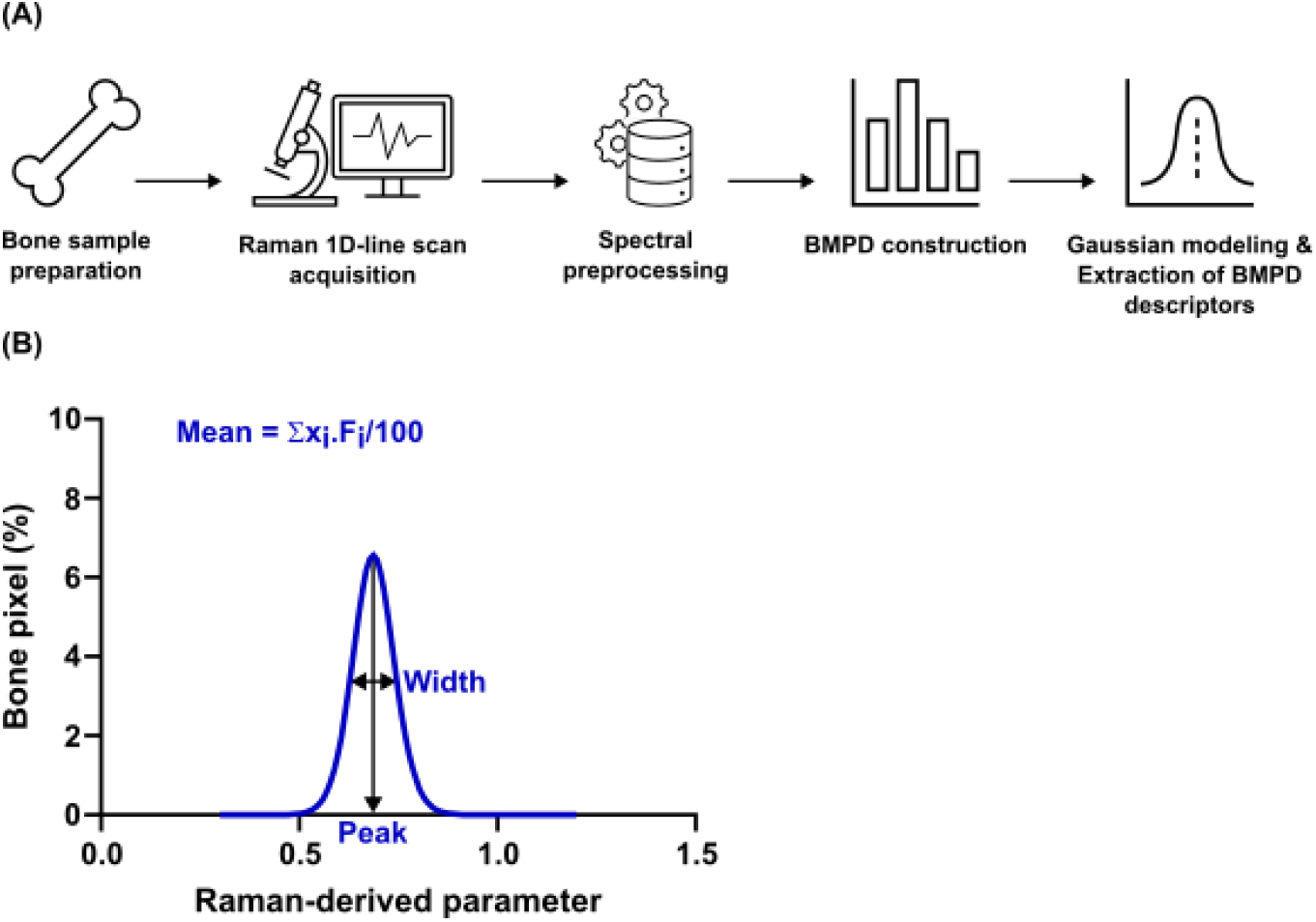
Overview of the quantitative Raman analysis (qRA) workflow for the construction of Bone Material Properties Distributions (BMPDs). (A) Bone specimens were prepared according to the selected histological processing method (decalcified paraffin embedding or undecalcified pMMA embedding) and analyzed by one-dimensional Raman line-scan acquisition across individual bone structural units. Raman spectra were subsequently subjected to standardized preprocessing, including baseline correction, normalization, smoothing, and embedding-medium subtraction. Raman-derived biochemical parameters were calculated for each bone pixel, and their pixel-wise distributions were represented as normalized histograms (BMPDs). (B) Each BMPD was then fitted with a Gaussian function to extract three quantitative descriptors: Mean, Peak, and Width, describing the central tendency, most probable value, and spatial heterogeneity of each bone material property.

**Figure 2.**
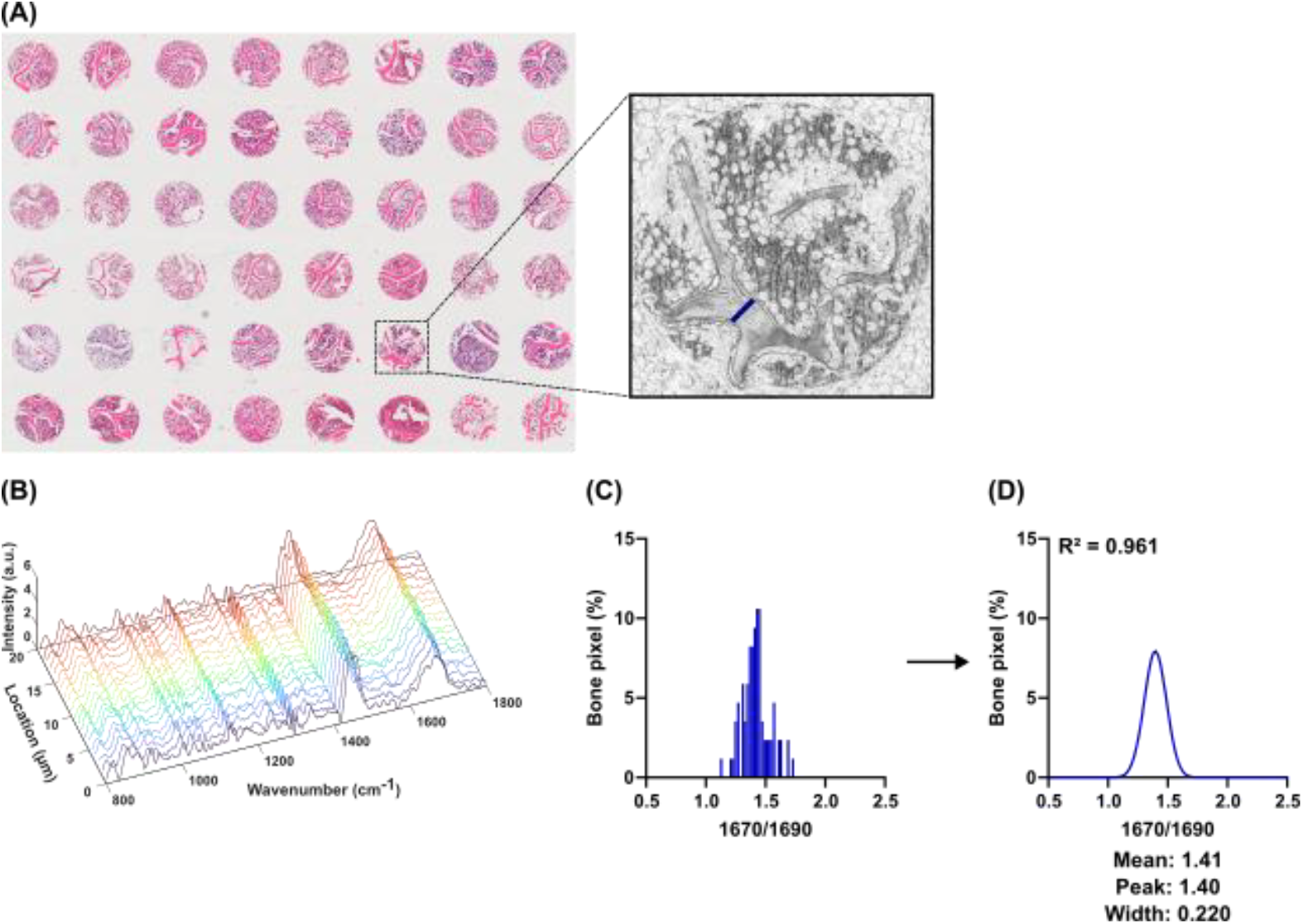
Construction of a BMPD from Raman line-scan acquisitions in paraffin-embedded bone tissue. (A) Hematoxylin-eosin-saffron (HES)-stained section of the BM481a tissue microarray (TMA) containing duplicate decalcified paraffin-embedded osteomedullary biopsies from 24 individuals without bone disease. The inset shows a corresponding core on an adjacent serial section mounted on a polished stainless-steel slide and used for Raman microspectroscopy. The blue line indicates the Raman line-scan acquisition used for BMPD construction. (B) Representative preprocessed Raman spectra corresponding to the first 20 acquisition points along the line scan. (C) BMPD of the collagen maturity parameter (1670/1690), generated from the pixel-wise Raman values measured along the line scan. (D) Gaussian modelling of the BMPD, providing the coefficient of determination (R^2^) and the three quantitative descriptors derived from the fitted distribution: 1670/1690_mean, 1670/1690_peak, and 1670/1690_width.

**Figure 3.**
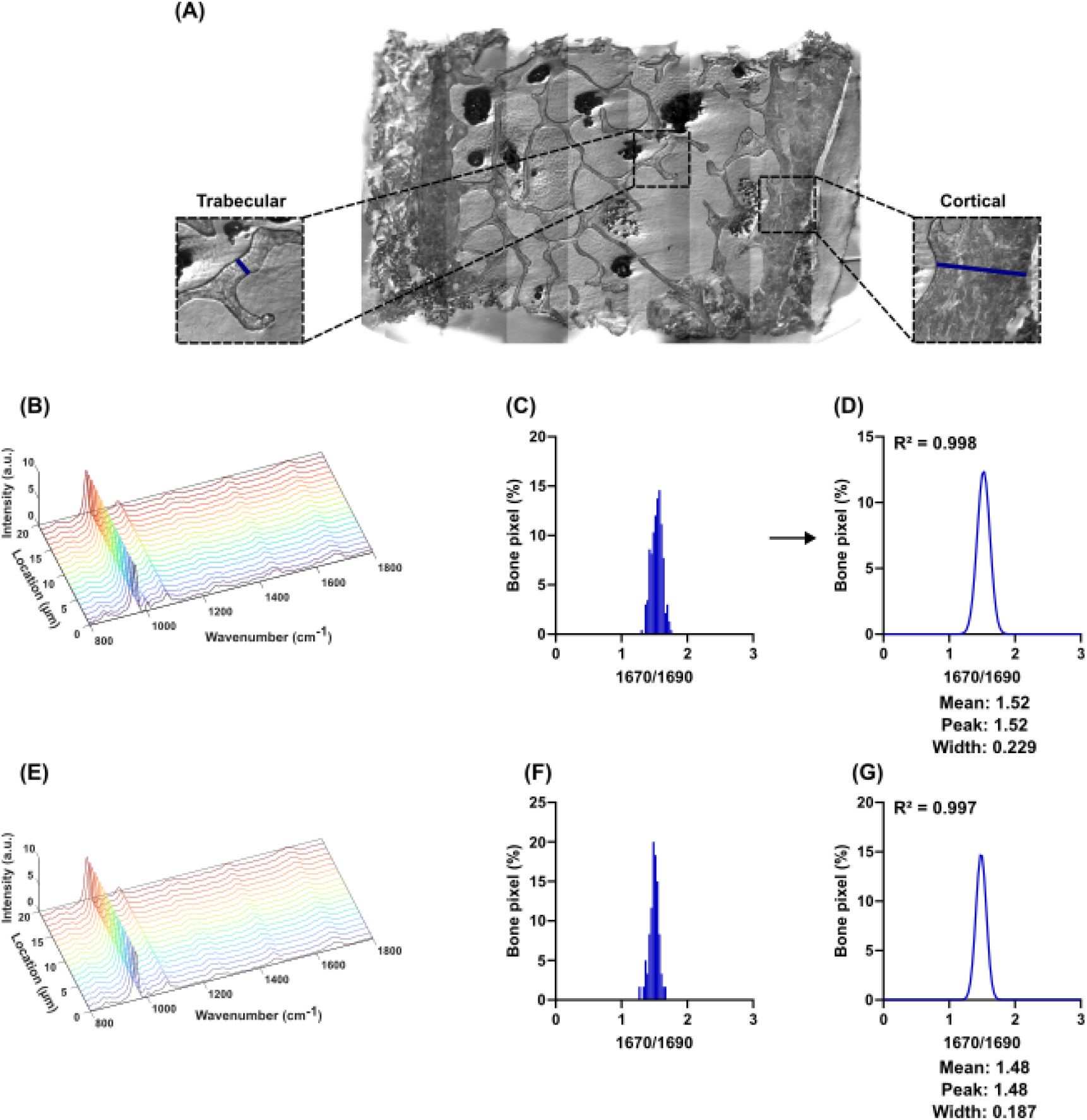
Construction of BMPDs from Raman line-scan acquisitions in pMMA-embedded bone tissue. (A) Reflected-light image of the surface of a pMMA-embedded bone biopsy as viewed with the Raman microspectrometer. The trabecular and cortical insets indicate the regions of interest selected for Raman microspectroscopy. Blue lines indicate the Raman line-scan acquisitions used for BMPD construction. (B) Representative preprocessed Raman spectra of cortical bone corresponding to the first 20 acquisition points along the line scan. (C) BMPD of the collagen maturity parameter (1670/1690) in cortical bone, generated from the pixel-wise Raman values measured along the line scan. (D) Gaussian modeling of the cortical BMPD, providing the coefficient of determination (R^2^) and the three quantitative descriptors derived from the fitted distribution: 1670/1690_mean, 1670/1690_peak, and 1670/1690_width. (E) Representative preprocessed Raman spectra of trabecular bone corresponding to the first 20 acquisition points along the line scan. (F) BMPD of the collagen maturity parameter (1670/1690) in trabecular bone, generated from the pixel-wise Raman values measured along the line scan. (G) Gaussian modeling of the trabecular BMPD, providing the coefficient of determination (R^2^) and the three quantitative descriptors derived from the fitted distribution: 1670/1690_mean, 1670/1690_peak, and 1670/1690_width.

Across all Raman-derived parameters and specimen preparation methods, BMPDs were predominantly unimodal. Gaussian functions provided an accurate approximation of the experimental distributions, although the goodness of fit varied slightly according to the biochemical parameter analyzed. Mean R^2^ values for all parameters and preparation methods are summarized in Supplementary Table S1.

### 3.2. Validation of line-scan Raman acquisition for quantitative biochemical assessment

To determine whether one-dimensional line-scan Raman acquisition provides quantitative biochemical information comparable to conventional two-dimensional ROI-based Raman mapping, Raman-derived BMPD descriptors obtained from both acquisition strategies were compared in paraffin-embedded decalcified bone biopsies. Excellent agreement was observed between line-scan and ROI-based acquisitions for both BMPD Mean and BMPD Width across all organic matrix parameters (Supplementary Table 2). Mean biases were negligible (−0.001 to 0.001), limits of agreement were narrow, and both Spearman’s correlation coefficients (ρ = 0.918–0.999) and ICC values (0.871–1.000) demonstrated excellent reproducibility of the reduced-dimensional acquisition strategy. BMPD_peak exhibited greater variability than BMPD_mean and BMPD_width. Although most peak-derived descriptors remained strongly correlated between acquisition modalities (ρ = 0.696–0.937), agreement was more heterogeneous (ICC = 0.730– 0.912). The PEN/CH_2_ peak parameter showed the lowest reproducibility (ICC = 0.385; 95% CI, −0.048 to 0.699), reflecting increased sensitivity of the peak descriptor to subtle changes in the fitted gaussian distribution. Replacing conventional ROI Raman mapping by a single line scan reduced the acquisition time from approximately 4,260 minutes to 42 minutes for an equivalent sampling area, representing an approximately 100-fold reduction while preserving the quantitative biochemical information required for BMPD analysis. Likewise, the robustness of the line-scan acquisition strategy was subsequently evaluated in undecalcified PMMA-embedded bone biopsies, allowing assessment of both organic matrix and mineral-related Raman parameters (Supplementary Table 3). Overall, agreement between 1D line-scan and 2D ROI acquisitions was excellent for BMPD_mean, with mean biases ranging from −0.003 to 0.003, narrow limits of agreement, Spearman’s correlation coefficients between 0.877 and 0.998, and ICC values ranging from 0.903 to 1.000. Similarly, BMPD_width showed excellent reproducibility for all parameters investigated. Mean biases remained negligible (−0.005 to 0.002), while correlation coefficients (ρ = 0.914–0.998) and ICC values (0.946– 1.000) confirmed that the line-scan approach accurately preserved the spatial heterogeneity of both the organic and mineral phases of bone tissue. As observed for paraffin-embedded specimens, BMPD_peak exhibited greater variability than the other gaussian-derived descriptors. Nevertheless, agreement remained excellent for the majority of Raman parameters, with ICC values ranging from 0.836 to 0.998. Lower reproducibility was observed for the crystallinity and collagen maturity (1670/1690 cm^−1)^ peak descriptors, which also displayed the lowest Spearman’s correlation coefficients (ρ = 0.514 and 0.595, respectively). Despite this increased variability, all remaining peak-derived parameters demonstrated excellent agreement between acquisition strategies.

### 3.3. Interindividual variability of Raman-derived bone material properties assessed by BMPDs

For each biochemical parameter, individual line scan BMPDs were generated for every patient included in the paraffin-embedded and pMMA-embedded cohorts. All individual BMPDs were plotted on a common graph to visualize the distribution and variability of Raman-derived bone material properties across specimens (Figures 4–6). The mean BMPD ± standard deviation was then calculated for each parameter and tissue preparation group. In addition, the mean ± standard deviation values of the BMPD descriptors (mean, peak, and width) were calculated separately for trabecular bone from the paraffin-embedded cohort, trabecular bone from the pMMA-embedded cohort, and cortical bone from the pMMA-embedded cohort. These quantitative descriptors are presented in Tables 1–3. Across all tissue preparations, BMPD_mean and BMPD_peak were highly consistent, with only small differences observed between the two descriptors for each biochemical parameter. In contrast, BMPD_width exhibited greater inter-individual variability, reflecting differences in the spatial heterogeneity of bone material properties among specimens. Parameters associated with 1670/1690, 1670/1640, Hyp/Pro, and mineral crystallinity displayed the lowest variability, whereas advanced glycation end-product (AGE)-related parameters (CML/CH_2_ and PEN/CH_2_) and GAG/Amide III showed comparatively larger dispersion. Comparison of trabecular and cortical bone in the pMMA cohort revealed generally similar BMPD_mean and BMPD_peak values for most Raman-derived parameters. However, cortical bone tended to exhibit narrower BMPDs, as reflected by lower BMPD_width values for most organic matrix parameters, indicating reduced spatial heterogeneity compared with trabecular bone. Mineral-related parameters showed limited differences between trabecular and cortical compartments, with crystallinity and carbonate substitution exhibiting particularly consistent BMPD descriptors. Together, these reference values establish normative BMPD descriptors for Raman-derived bone material parameters in both decalcified paraffin-embedded and undecalcified pMMA-embedded human bone biopsies, providing quantitative benchmarks for future studies investigating alterations in bone material composition and heterogeneity.

**Table 1.** BMPD descriptors in paraffin-embedded trabecular bone. BMPD-derived parameters (mean, peak and width) are reported for Raman-derived biochemical indices measured in decalcified paraffin-embedded bone biopsies. These values represent reference BMPD descriptors of decalcified human osteomedullary biopsies embedded in paraffin. Data are presented as mean ± standard deviation across individuals in the reference cohort.

|  | Mean | Peak | Width |
| --- | --- | --- | --- |
| 1670/1690 | 1.441 ± 0.026 | 1.427 ± 0.029 | 0.250 ± 0.041 |
| 1670/1640 | 1.498 ± 0.018 | 1.481 ± 0.019 | 0.289 ± 0.044 |
| Hyp/Pro | 0.839 ± 0.035 | 0.820 ± 0.035 | 0.208 ± 0.033 |
| CML/CH<sub>2</sub> | 0.178 ± 0.060 | 0.154 ± 0.050 | 0.156 ± 0.041 |
| PEN/CH<sub>2</sub> | 0.053 ± 0.018 | 0.031 ± 0.008 | 0.078 ± 0.043 |
| GAG/Amide III | 0.735 ± 0.112 | 0.662 ± 0.106 | 0.455 ± 0.118 |

**Table 2.** BMPD descriptors in undecalcified pMMA-embedded trabecular bone. BMPD-derived parameters (mean, peak and width) are reported for Raman-derived biochemical indices measured in undecalcified human transiliac bone biopsies embedded in pMMA. Data are presented as mean ± standard deviation across individuals in the PMMA cohort.

|  | Mean | Peak | Width |
| --- | --- | --- | --- |
| 1670/1690 | 1.507 ± 0.029 | 1.503 ± 0.032 | 0.262 ± 0.065 |
| 1670/1640 | 1.453 ± 0.054 | 1.430 ± 0.070 | 0.236 ± 0.083 |
| Hyp/Pro | 0.710 ± 0.023 | 0.686 ± 0.033 | 0.130 ± 0.072 |
| CML/CH<sub>2</sub> | 0.255 ± 0.034 | 0.237 ± 0.036 | 0.121 ± 0.053 |
| PEN/CH<sub>2</sub> | 0.313 ± 0.049 | 0.263 ± 0.046 | 0.128 ± 0.076 |
| GAG/Amide III | 0.462 ± 0.038 | 0.421 ± 0.039 | 0.154 ± 0.084 |
| v1PO<sub>4</sub>/Amide I | 7.32 ± 0.67 | 7.18 ± 0.96 | 1.72 ± 0.92 |
| v1PO<sub>4</sub>/Amide III | 9.69 ± 0.91 | 9.85 ± 1.06 | 2.42 ± 1.08 |
| v1PO<sub>4</sub>/CH<sub>2</sub> | 7.98 ± 0.71 | 8.06 ± 1.44 | 2.45 ± 1.15 |
| v1PO<sub>4</sub>/Hyp+Pro | 5.84 ± 0.54 | 5.89 ± 0.73 | 1.79 ± 0.72 |
| Crystallinity | 0.0572 ± 0.0005 | 0.0571 ± 0.0005 | 0.0017 ± 0.0004 |
| v1CO<sub>3</sub>/v1PO<sub>4</sub> | 0.199 ± 0.012 | 0.197 ± 0.011 | 0.027 ± 0.007 |

**Table 3.**
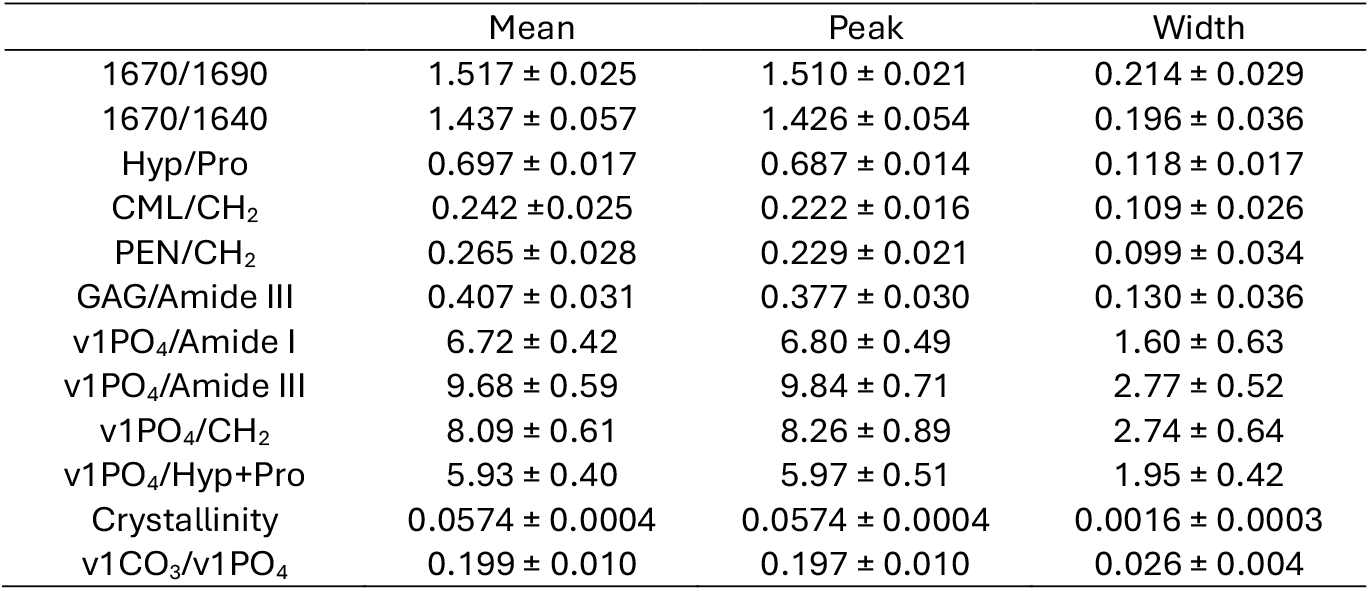
BMPD descriptors in undecalcified pMMA-embedded cortical bone. BMPD-derived parameters (mean, peak and width) are reported for Raman-derived biochemical indices measured in undecalcified human transiliac bone biopsies embedded in pMMA. Data are presented as mean ± standard deviation across individuals in the PMMA cohort.

**Figure 4.**
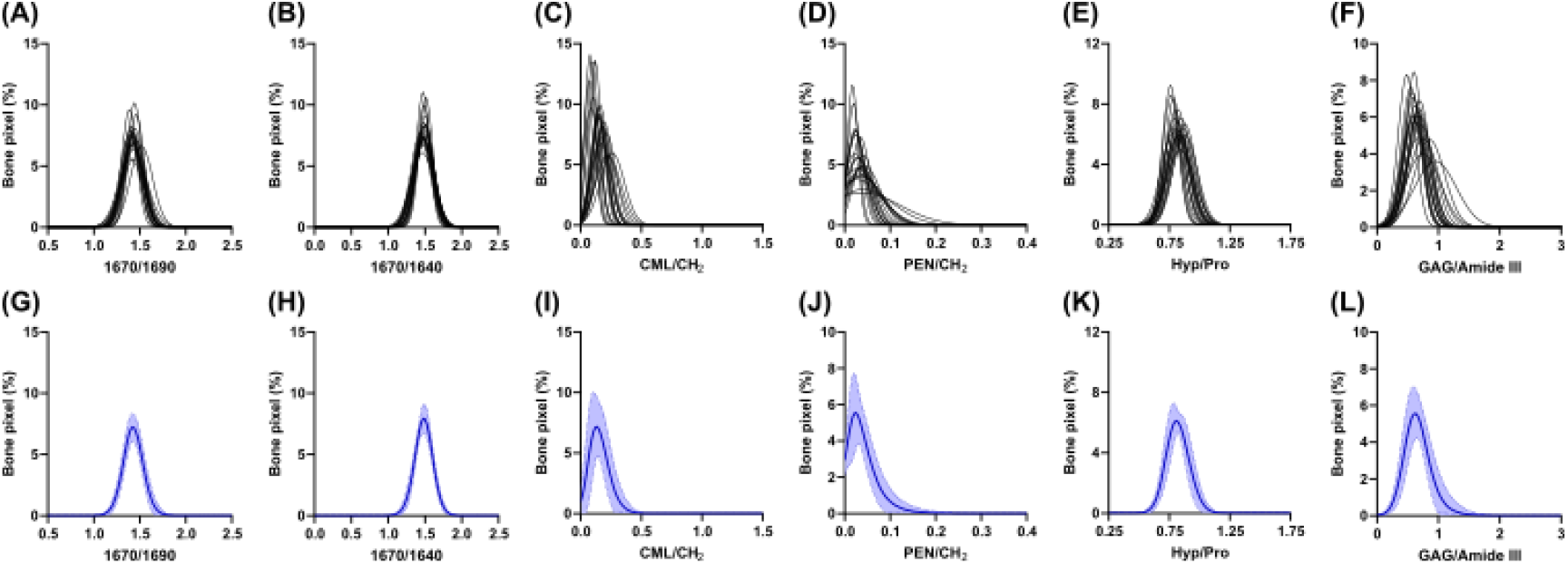
BMPDs of Raman-derived biochemical parameters in trabecular bone from paraffin-embedded specimens. (A– F) Individual BMPDs obtained from the 21 decalcified paraffin-embedded osteomedullary biopsies. Each curve represents the normalized distribution of a single patient for (A) 1670/1690 cm^−1^, (B) 1670/1640 cm^−1^, (C) CML/CH_2_, (D) PEN/CH_2_, (E) Hyp/Pro, and (F) GAG/Amide III ratios. (G–L) Corresponding mean BMPDs calculated from the 21 individual distributions for (G) 1670/1690 cm^−1^, (H) 1670/1640 cm^−1^, (I) CML/CH_2_, (J) PEN/CH_2_, (K) Hyp/Pro, and (L) GAG/Amide III ratios. Solid lines represent the mean BMPD, and shaded areas indicate the standard deviation (SD) at each bin.

**Figure 5.**
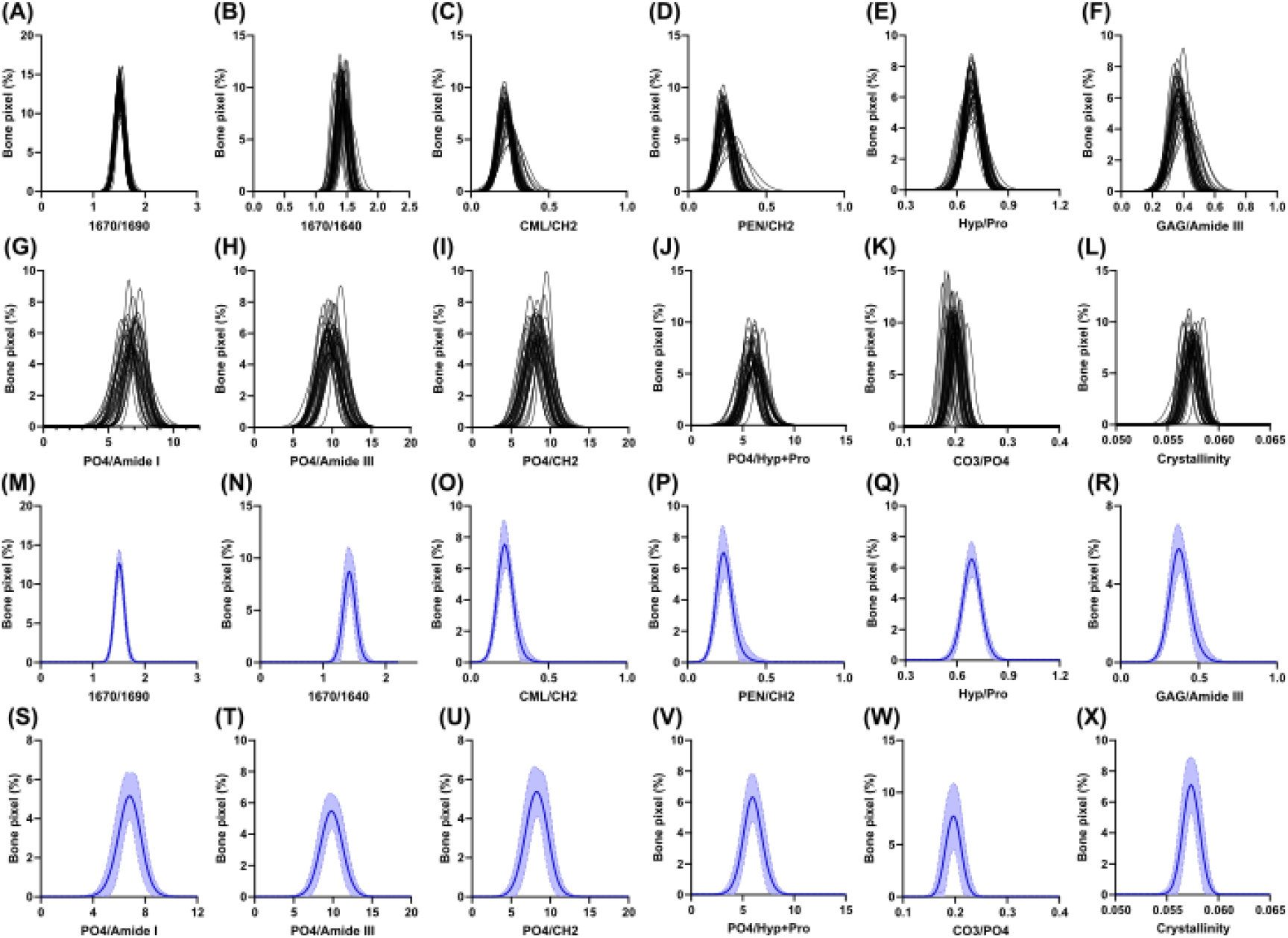
BMPDs of Raman-derived biochemical parameters in cortical bone from pMMA-embedded specimens. (A–F) Individual BMPDs obtained from the 38 undecalcified pMMA-embedded iliac crest biopsies. Each curve represents the normalized distribution of a single patient for (A) 1670/1690 cm^−1^, (B) 1670/1640 cm^−1^, (C) CML/CH_2_, (D) PEN/CH_2_, (E) Hyp/Pro, (F) GAG/Amide III, (G) PO_4_/Amide I, (H) PO_4_/Amide III, (I) PO_4_/CH_2_, (J) PO_4_/Hyp+Pro, (K) CO_3_/PO_4_ ratios and (L) Crystallinity. (M– X) Corresponding mean BMPDs calculated from the 38 individual distributions for (M) 1670/1690 cm^−1^, (N) 1670/1640 cm^−1^, (O) CML/CH_2_, (P) PEN/CH_2_, (Q) Hyp/Pro, (R) GAG/Amide III, (S) PO_4_/Amide I, (T) PO_4_/Amide III, (U) PO_4_/CH_2_, (V) PO_4_/Hyp+Pro, (W) CO_3_/PO_4_ ratios and (X) Crystallinity. Solid lines represent the mean BMPD, and shaded areas indicate the standard deviation (SD) at each bin.

**Figure 6.**
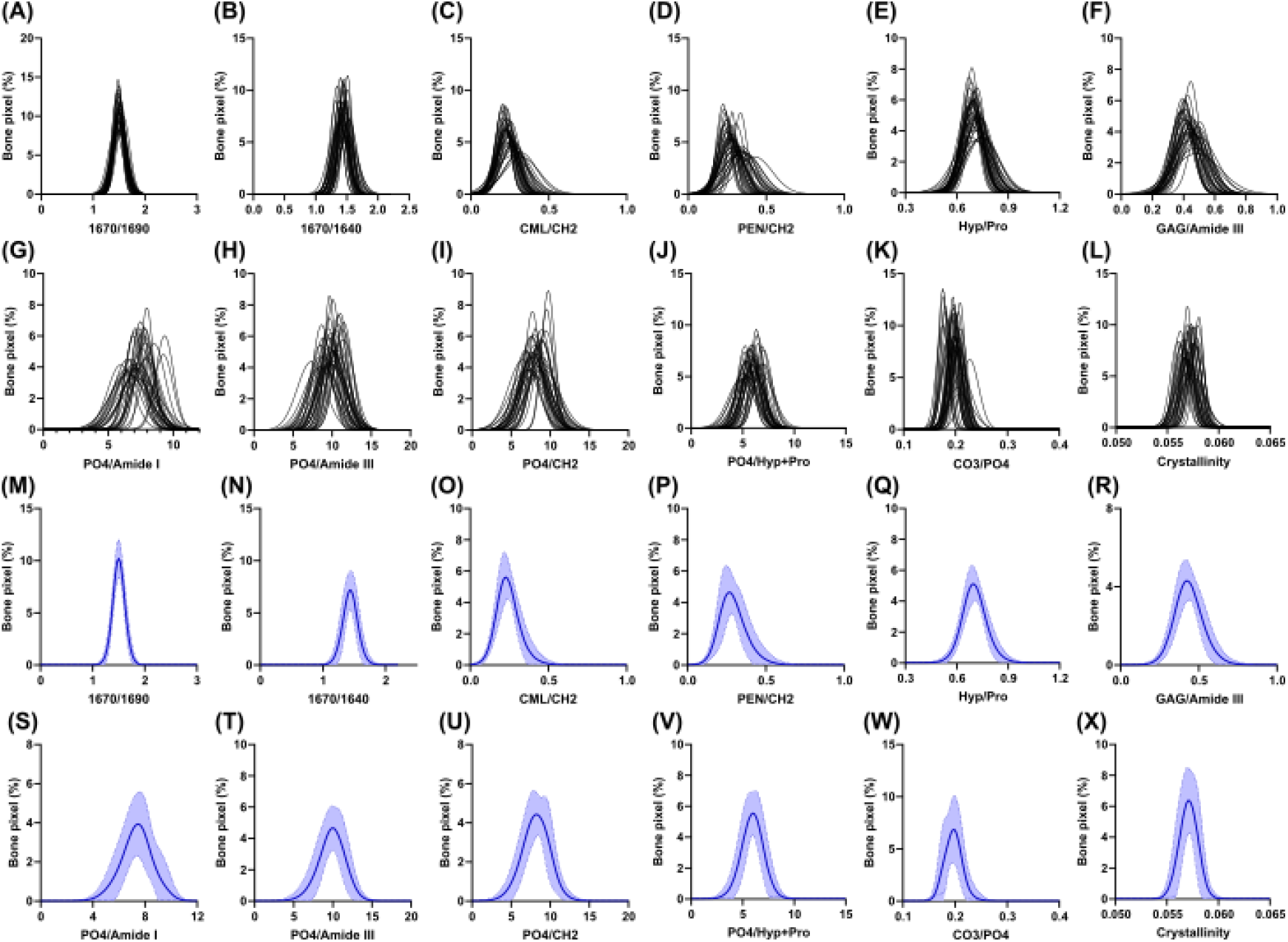
BMPDs of Raman-derived biochemical parameters in trabecular bone from pMMA-embedded specimens. (A– F) Individual BMPDs obtained from the 38 undecalcified pMMA-embedded iliac crest biopsies. Each curve represents the normalized distribution of a single patient for (A) 1670/1690 cm^−1^, (B) 1670/1640 cm^−1^, (C) CML/CH_2_, (D) PEN/CH_2_, (E) Hyp/Pro, (F) GAG/Amide III, (G) PO_4_/Amide I, (H) PO_4_/Amide III, (I) PO_4_/CH_2_, (J) PO_4_/Hyp+Pro, (K) CO_3_/PO_4_ ratios and (L) Crystallinity. (M– X) Corresponding mean BMPDs calculated from the 38 individual distributions for (M) 1670/1690 cm^−1^, (N) 1670/1640 cm^−1^, (O) CML/CH_2_, (P) PEN/CH_2_, (Q) Hyp/Pro, (R) GAG/Amide III, (S) PO_4_/Amide I, (T) PO_4_/Amide III, (U) PO_4_/CH_2_, (V) PO_4_/Hyp+Pro, (W) CO_3_/PO_4_ ratios and (X) Crystallinity. Solid lines represent the mean BMPD, and shaded areas indicate the standard deviation (SD) at each bin.

The reproducibility of the BMPD-derived descriptors was evaluated by comparing technical and biological coefficients of variation (CVs) (Table 4). Technical variability was determined from five repeated acquisitions of the same Raman line scan, with complete instrument recalibration, including spectral calibration using the internal silicon standard and laser beam realignment, performed before each acquisition. Interindividual biological variability was assessed across the two cohorts. Across all Raman-derived biochemical parameters, technical CVs were consistently lower than interindividual biological CVs, demonstrating the high reproducibility of the BMPD methodology. Technical variability was generally lower for pMMA-embedded specimens than for paraffin-embedded specimens, with technical CVs typically below 2% for BMPD_mean and BMPD_peak and almost below 4% for BMPD_width in pMMA-embedded bone. In comparison, repeated measurements in paraffin-embedded specimens showed slightly higher technical variability, particularly for BMPD_width and AGE-related parameters. Interindividual biological variability differed substantially among biochemical parameters. Among the organic matrix parameters, the 1670/1690, 1670/1640, and Hyp/Pro ratios exhibited relatively low interindividual biological CVs for BMPD_mean and BMPD_peak, whereas CML/CH_2_ and PEN/CH_2_ ratios displayed markedly higher inter-individual variability. GAG/Amide III showed intermediate variability. For mineral-related parameters, assessed only in pMMA-embedded specimens, crystallinity exhibited the lowest biological variability of all parameters (CV ≤1% for BMPD_mean and BMPD_peak), whereas mineral-to-matrix ratios and carbonate substitution displayed moderate biological variability (approximately 5–13%). In contrast, BMPD_width exhibited substantially larger biological CVs for all mineral parameters, particularly in trabecular bone. Regardless of tissue preparation or biochemical parameter, BMPD_width consistently displayed greater biological variability than BMPD_mean or BMPD_peak, indicating that the spatial heterogeneity of bone material properties varied more markedly between individuals than their average or most prevalent values.

**Table 4.** Coefficients of variation (CVs) of Raman-derived BMPD descriptors. CVs were calculated to assess both technical and biological variabilities of BMPD-derived descriptors. Technical CVs were determined by five repeated Raman acquisitions of the same line scan, with complete instrument recalibration between acquisitions, including spectral calibration using the internal silicon standard and laser beam realignment prior to each measurement. BMPDs were reconstructed after each acquisition, and the corresponding descriptors were calculated. Biological CVs were calculated across individuals within each cohort (21 decalcified paraffin-embedded biopsies and 38 undecalcified pMMA-embedded biopsies). Results are presented separately for trabecular bone from decalcified paraffin-embedded biopsies, trabecular bone from undecalcified pMMA-embedded biopsies, and cortical bone from undecalcified pMMA-embedded biopsies.

| Parameters | Paraffin-embedded trabecular bone |  | pMMA-embedded trabecular bone |  | pMMA-embedded cortical bone |  |
| --- | --- | --- | --- | --- | --- | --- |
| Variability | Technical | Biological | Technical | Biological | Technical | Biological |
| Mean |  |  |  |  |  |  |
| 1670/1690 | 0.4 | 1.8 | 0.4 | 1.9 | 0.6 | 1.6 |
| 1670/1640 | 1.2 | 1.2 | 0.2 | 3.7 | 0.1 | 3.9 |
| Hyp/Pro | 1.1 | 4.2 | 0.3 | 3.2 | 0.2 | 2.4 |
| CML/CH<sub>2</sub> | 3.3 | 33.7 | 1.1 | 13.4 | 1.0 | 10.1 |
| PEN/CH<sub>2</sub> | 0.6 | 34.0 | 0.1 | 15.5 | 1.5 | 10.7 |
| GAG/Amide III | 1.9 | 15.2 | 0.8 | 8.2 | 0.1 | 7.7 |
| v1PO<sub>4</sub>/Amide I | - | - | 0.7 | 9.1 | 0.2 | 6.2 |
| v1PO<sub>4</sub>/Amide III | - | - | 1.4 | 9.4 | 0.2 | 6.1 |
| v1PO<sub>4</sub>/CH<sub>2</sub> | - | - | 1.4 | 8.9 | 0.1 | 7.6 |
| v1PO<sub>4</sub>/Hyp+Pro | - | - | 1.2 | 9.2 | 0.3 | 6.7 |
| Crystallinity | - | - | 0.2 | 1.0 | 0.0 | 0.8 |
| v1CO<sub>3</sub>/v1PO<sub>4</sub> | - | - | 0.8 | 5.9 | 0.1 | 5.1 |
| Peak |  |  |  |  |  |  |
| 1670/1690 | 0.7 | 2.0 | 0.3 | 2.2 | 0.6 | 1.4 |
| 1670/1640 | 1.3 | 1.3 | 0.2 | 4.9 | 0.1 | 3.8 |
| Hyp/Pro | 1.2 | 4.3 | 0.1 | 4.9 | 0.3 | 2.0 |
| CML/CH<sub>2</sub> | 3.9 | 32.5 | 2.0 | 15.3 | 0.6 | 7.4 |
| PEN/CH<sub>2</sub> | 6.5 | 25.8 | 0.9 | 17.5 | 0.6 | 9.0 |
| GAG/Amide III | 5.0 | 16.0 | 0.8 | 9.3 | 0.5 | 7.8 |
| v1PO<sub>4</sub>/Amide I | - | - | 0.8 | 13.4 | 0.2 | 7.2 |
| v1PO<sub>4</sub>/Amide III | - | - | 1.7 | 10.7 | 0.1 | 7.2 |
| v1PO<sub>4</sub>/CH<sub>2</sub> | - | - | 1.7 | 17.9 | 0.1 | 10.8 |
| v1PO<sub>4</sub>/Hyp+Pro | - | - | 1.6 | 12.4 | 0.3 | 8.6 |
| Crystallinity | - | - | 0.2 | 1.0 | 0.0 | 0.8 |
| v1CO<sub>3</sub>/v1PO<sub>4</sub> | - | - | 0.8 | 5.6 | 0.1 | 5.1 |
| Width |  |  |  |  |  |  |
| 1670/1690 | 2.7 | 16.4 | 0.8 | 24.8 | 1.1 | 13.4 |
| 1670/1640 | 3.0 | 15.2 | 1.7 | 35.2 | 1.0 | 18.5 |
| Hyp/Pro | 4.2 | 15.9 | 1.0 | 55.3 | 3.4 | 14.5 |
| CML/CH<sub>2</sub> | 4.8 | 26.3 | 1.4 | 43.7 | 4.7 | 24.2 |
| PEN/CH<sub>2</sub> | 6.4 | 55.1 | 0.4 | 59.0 | 1.9 | 34.0 |
| GAG/Amide III | 4.3 | 25.9 | 0.9 | 54.6 | 3.6 | 27.5 |
| v1PO<sub>4</sub>/Amide I | - | - | 2.5 | 53.6 | 1.8 | 39.4 |
| v1PO<sub>4</sub>/Amide III | - | - | 0.5 | 44.7 | 0.8 | 18.6 |
| v1PO<sub>4</sub>/CH<sub>2</sub> | - | - | 0.3 | 46.9 | 0.5 | 23.4 |
| v1PO<sub>4</sub>/Hyp+Pro | - | - | 0.6 | 40.1 | 0.3 | 21.4 |
| Crystallinity | - | - | 4.1 | 22.9 | 2.1 | 17.2 |
| v1CO<sub>3</sub>/v1PO<sub>4</sub> | - | - | 3.9 | 24.7 | 1.7 | 15.2 |

Interindividual biological variability was also generally lower in cortical than in trabecular bone for both organic and mineral matrix parameters. In order to evaluate whether decalcification routinely done in pathology lab alter the BMPD descriptors, we compared the 18 BMPD descriptors in decalcified paraffin-embedded osteomedullary biopsies and EDTA-decalcified pMMA-embedded biopsies. As shown in Supplementary Figure S4, none of the investigated BMPD descriptor distributions were significantly different between the two bone sample cohorts, suggesting that the rapid decalcification process used in pathology lab does not alter significantly the BMPD descriptors.

### 3.4. Representativeness of a single trabecular line scan

To determine whether a single Raman line scan was representative of the bone material properties of an individual biopsy, variance component analyses were performed using linear mixed-effects models on three independent trabeculae from each biopsy. Between-patient (τ^2^) and within-patient (σ^2^) variance components, together with the corresponding single-measure intraclass correlation coefficients (ICC(1)), are summarized in Table 5. For BMPD_mean, ICC values were consistently high across all Raman-derived biochemical parameters, ranging from 0.90 to 0.99 in both paraffin-embedded and pMMA-embedded specimens. Comparable ICC values were obtained in trabecular and cortical bone, indicating that inter-individual differences accounted for most of the observed variability, whereas variability among trabeculae within the same biopsy remained limited.

**Table 5.** Variance component analysis, intraclass correlation coefficients (ICC), and estimated number of bone structural units required for reliable BMPD measurements. Variance components were estimated using linear mixed-effects models based on three independent trabecular/cortical Raman line scans obtained from each biopsy. Between-patient (τ^2^) and within-patient (σ^2^) variance components, single-measure intraclass correlation coefficients (ICC(1)) with their 95% confidence intervals, and the estimated number of independent Raman line scans required to achieve a target reliability of ICC ≥ 0.80 are reported. The required number of line scans was calculated using the Spearman–Brown prophecy formula.

| | $\tau^2$ | | $\sigma^2$ | | ICC(1) | | 95% CI | | # of Raman line scans | |
| --- | --- | --- | --- | --- | --- | --- | --- | --- | --- | --- |
|  | Trab. | Cort. | Trab. | Cort. | Trab. | Cort. | Trab. | Cort. | Trab. | Cort. |
| **BMPD<sub>mean</sub>** |  |  |  |  |  |  |  |  |  |  |
| **pMMA-embedded** |  |  |  |  |  |  |  |  |  |  |
| 1670/1690 | 0.00184 | 0.00048 | 0.00013 | 0.00005 | 0.934 | 0.901 | 0.832-0.969 | 0.762-0.968 | 1 | 1 |
| 1670/1640 | 0.00164 | 0.00201 | 0.00014 | 0.00012 | 0.923 | 0.944 | 0.807-0.964 | 0.858-0.973 | 1 | 1 |
| Hyp/Pro | 0.00107 | 0.00040 | 0.00010 | 0.00001 | 0.915 | 0.969 | 0.789-0.960 | 0.920-0.986 | 1 | 1 |
| CML/CH<sub>2</sub> | 0.00320 | 0.00193 | 0.00031 | 0.00010 | 0.911 | 0.952 | 0.778-0.958 | 0.877-0.977 | 1 | 1 |
| PEN/CH<sub>2</sub> | 0.00659 | 0.00125 | 0.00041 | 0.00002 | 0.942 | 0.987 | 0.849-0.973 | 0.966-0.994 | 1 | 1 |
| GAG/Amide III | 0.00759 | 0.00347 | 0.00057 | 0.00003 | 0.930 | 0.991 | 0.823-0.967 | 0.976-0.996 | 1 | 1 |
| v1PO<sub>4</sub>/Amide I | 0.68572 | 0.30433 | 0.02133 | 0.02262 | 0.970 | 0.931 | 0.920-0.986 | 0.827-0.967 | 1 | 1 |
| v1PO<sub>4</sub>/Amide III | 1.60394 | 0.46923 | 0.04090 | 0.00114 | 0.975 | 0.998 | 0.934-0.989 | 0.993-0.999 | 1 | 1 |
| v1PO<sub>4</sub>/CH<sub>2</sub> | 0.98161 | 0.35913 | 0.02739 | 0.00195 | 0.973 | 0.995 | 0.928-0.988 | 0.985-0.998 | 1 | 1 |
| v1PO<sub>4</sub>/Hyp+Pro | 0.69740 | 0.24006 | 0.01120 | 0.00173 | 0.984 | 0.993 | 0.958-0.993 | 0.981-0.997 | 1 | 1 |
| Crystallinity | 0.00000 | 0.00000 | 0.00000 | 0.00000 | 0.901 | 0.877 | 0.757-0.954 | 0.708-0.938 | 1 | 1 |
| v1CO<sub>3</sub>/v1PO<sub>4</sub> | 0.00038 | 0.00010 | 0.00003 | 0.00001 | 0.927 | 0.907 | 0.815-0.966 | 0.772-0.955 | 1 | 1 |
| **Paraffin-embedded** |  |  |  |  |  |  |  |  |  |  |
| 1670/1690 | 0.00115 | - | 0.00012 | - | 0.904 | - | 0.767-0.954 | - | 1 | - |
| 1670/1640 | 0.00271 | - | 0.00014 | - | 0.949 | - | 0.871-0.976 | - | 1 | - |
| Hyp/Pro | 0.00056 | - | 0.00001 | - | 0.976 | - | 0.937-0.989 | - | 1 | - |
| CML/CH<sub>2</sub> | 0.00336 | - | 0.00011 | - | 0.969 | - | 0.918-0.986 | - | 1 | - |
| PEN/CH<sub>2</sub> | 0.00001 | - | 0.00000 | - | 0.947 | - | 0.864-0.975 | - | 1 | - |
| GAG/Amide III | 0.00074 | - | 0.00001 | - | 0.990 | - | 0.973-0.995 | - | 1 | - |
| **BMPD<sub>width</sub>** |  |  |  |  |  |  |  |  |  |  |
| **pMMA-embedded** |  |  |  |  |  |  |  |  |  |  |
| 1670/1690 | 0.00560 | 0.00334 | 0.00015 | 0.00010 | 0.974 | 0.970 | 0.931-0.988 | 0.921-0.987 | 1 | 1 |
| 1670/1640 | 0.00340 | 0.00114 | 0.00009 | 0.00009 | 0.974 | 0.929 | 0.931-0.988 | 0.819-0.967 | 1 | 1 |
| Hyp/Pro | 0.00162 | 0.00383 | 0.00011 | 0.00004 | 0.935 | 0.990 | 0.834-0.970 | 0.974-0.996 | 1 | 1 |
| CML/CH<sub>2</sub> | 0.04914 | 0.00036 | 0.00064 | 0.00005 | 0.987 | 0.877 | 0.966-0.994 | 0.702-0.941 | 1 | 1 |
| PEN/CH<sub>2</sub> | 0.07121 | 0.00153 | 0.00042 | 0.00012 | 0.994 | 0.925 | 0.984-0.997 | 0.810-0.965 | 1 | 1 |
| GAG/Amide III | 0.04290 | 0.00121 | 0.00026 | 0.00003 | 0.994 | 0.978 | 0.984-0.997 | 0.941-0.990 | 1 | 1 |
| v1PO<sub>4</sub>/Amide I | 0.14485 | 0.32620 | 0.01081 | 0.00430 | 0.931 | 0.987 | 0.823-0.968 | 0.965-0.994 | 1 | 1 |
| v1PO<sub>4</sub>/Amide III | 0.57561 | 0.31472 | 0.02327 | 0.00842 | 0.961 | 0.974 | 0.896-0.982 | 0.931-0.988 | 1 | 1 |
| v1PO<sub>4</sub>/CH<sub>2</sub> | 0.40534 | 0.50291 | 0.02129 | 0.00738 | 0.950 | 0.986 | 0.869-0.977 | 0.962-0.994 | 1 | 1 |
| v1PO<sub>4</sub>/Hyp+Pro | 0.34370 | 0.27690 | 0.01219 | 0.01017 | 0.966 | 0.965 | 0.909-0.984 | 0.906-0.984 | 1 | 1 |
| Crystallinity | 0.00000 | 0.00000 | 0.00000 | 0.00000 | 0.929 | 0.903 | 0.820-0.967 | 0.760-0.954 | 1 | 1 |
| v1CO<sub>3</sub>/v1PO<sub>4</sub> | 0.00012 | 0.00007 | 0.00001 | 0.00000 | 0.904 | 0.963 | 0.764-0.855 | 0.902-0.983 | 1 | 1 |
| Paraffin-embedded |  |  |  |  |  |  |  |  |  |  |
| 1670/1690 | 0.00022 | - | 0.00005 | - | 0.815 | - | 0.585-0.910 | - | 1 | - |
| 1670/1640 | 0.00009 | - | 0.00002 | - | 0.833 | - | 0.618-0.918 | - | 1 | - |
| Hyp/Pro | 0.00004 | - | 0.00001 | - | 0.860 | - | 0.671-0.932 | - | 1 | - |
| CML/CH<sub>2</sub> | 0.00214 | - | 0.00009 | - | 0.959 | - | 0.892-0.981 | - | 1 | - |
| PEN/CH<sub>2</sub> | 0.00001 | - | 0.00000 | - | 0.742 | - | 0.449-0.873 | - | 2 | - |
| GAG/Amide III | 0.04060 | - | 0.00061 | - | 0.985 | - | 0.960-0.993 | - | 1 | - |

A similar pattern was observed for BMPD_width, despite the greater biological variability previously observed for this descriptor. ICC values also exceeded 0.90 for nearly all Raman-derived biochemical parameters, indicating that differences in BMPD_width were predominantly attributable to inter-individual variability rather than to variability among trabeculae within the same biopsy. Thus, although the degree of bone material heterogeneity differed substantially between individuals, it remained consistent across independently analyzed trabeculae from the same biopsy. The only exception was the PEN/CH_2_ BMPD, which exhibited a lower ICC (0.74) in paraffin-embedded trabecular bone, reflecting greater within-biopsy variability. Application of the Spearman– Brown prophecy formula indicated that a single full-thickness Raman line scan spanning an entire trabecula or cortical wall was sufficient to achieve a reliability of ICC ≥ 0.80 for all Raman-derived biochemical parameters, except for PEN/CH_2_ BMPD_width in paraffin-embedded trabecular bone, for which two independent line scans would be required.

### 3.5. Independence of BMPD descriptors from demographic characteristics and illustrative application to pathological bone samples

To assess whether the proposed reference BMPD descriptors were influenced by demographic characteristics, linear regression analyses were performed for each Raman-derived biochemical parameter and each BMPD descriptor (BMPD_mean, BMPD_peak, and BMPD_width), using age and sex as explanatory variables. The corresponding false discovery rate (FDR)-adjusted p-values are summarized in the heatmap shown in Figure 7A. No significant association was identified between any BMPD descriptor and either age or sex after correction for multiple testing (FDR > 0.05 for all comparisons), indicating that the reference BMPD distributions were not measurably influenced by these demographic variables within the investigated cohorts.

**Figure 7.**
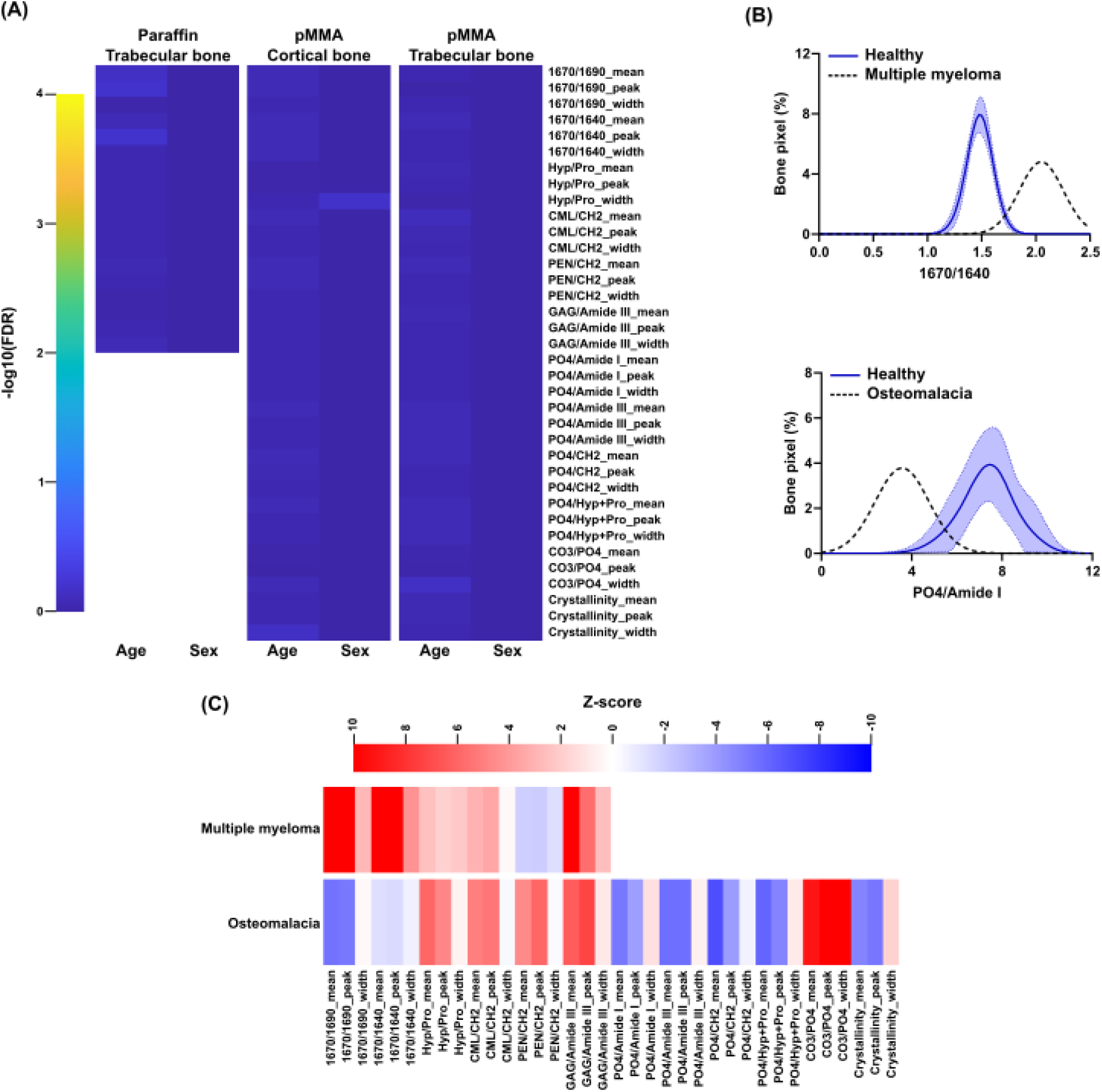
Independence of BMPD descriptors from age and sex and illustrative application of the BMPD reference distributions to pathological bone samples. (A) Heatmap summarizing the association between patient demographic characteristics and BMPD descriptors in the reference cohorts. Color intensity represents the −log_10_ (false discovery rate, FDR) obtained from multiple linear regression analyses testing the association of each BMPD descriptor (BMPD_mean, BMPD_peak, and BMPD_width for all Raman-derived biochemical parameters) with age or sex. No association remained statistically significant after Benjamini–Hochberg correction (FDR > 0.05 for all comparisons), indicating that BMPD descriptors are independent of age and sex in the reference populations. (B) Representative comparison of individual pathological BMPDs with the corresponding reference distributions. Top, 1670/1640 cm^−1^ BMPD from a decalcified paraffin-embedded bone marrow biopsy obtained from a patient with multiple myeloma (black dashed line), superimposed on the mean ± SD reference BMPD of the paraffin-embedded cohort (blue shaded area). Bottom, PO_4_/Amide I BMPD from an undecalcified pMMA-embedded transiliac bone biopsy obtained from a patient with osteomalacia (black dashed line), superimposed on the corresponding mean ± SD reference BMPD of the pMMA-embedded trabecular cohort (blue shaded area). These examples illustrate the visual comparison of an individual BMPD with the reference distribution. (C) Heatmap of standardized deviations (Z-scores) for the pathological cases shown in panel B. Z-scores were calculated for each BMPD descriptor (BMPD_mean, BMPD_peak, and BMPD_width) relative to the corresponding reference cohort (paraffin-embedded or pMMA-embedded). Positive and negative Z-scores indicate increases or decreases, respectively, relative to the reference population, allowing comprehensive visualization of biochemical alterations across all Raman-derived bone material parameters.

To illustrate the potential application of the reference BMPD distributions, representative pathological bone biopsies were compared with their corresponding reference cohorts. In the paraffin-embedded cohort, the BMPD of the 1670/1640 ratio obtained from a patient with multiple myeloma displayed a rightward shift relative to the reference distribution (Figure 7B). Likewise, in the pMMA-embedded cohort, the trabecular BMPD of the PO_4_/Amide I ratio obtained from a patient with osteomalacia displayed a leftward shift relative to the reference distribution (Figure 7B). These examples illustrate how individual BMPDs can be compared with the expected reference distributions to identify alterations in the spatial distribution of bone material properties. To facilitate the multidimensional interpretation of individual cases, BMPD descriptors were subsequently expressed as standardized deviations (Z-scores) relative to the corresponding reference cohort. The resulting heatmaps (Figure 7C) provide a comprehensive overview of the deviation of each pathological specimen from the reference population across all Raman-derived biochemical parameters and BMPD descriptors. Together, these results demonstrate that BMPDs constitute not only a quantitative framework for establishing reference distributions of Raman-derived bone material properties but also a practical tool for identifying and visualizing deviations from normal bone material composition in individual pathological bone biopsies.

## 4. Discussion

In the present study, we introduce a qRA framework that extends the concept of tissue composition assessment beyond conventional mean Raman values by generating BMPDs. Inspired by the BMDD approach established for qBEI, BMPDs describe not only the average value of Raman-derived biochemical parameters but also their spatial distribution and heterogeneity across successive BSUs. Using human reference bone biopsies prepared according to two widely used histological protocols, we demonstrate that these distributions are consistently approximated by gaussian functions, can be summarized using a limited number of robust descriptors, and are highly reproducible across tissue preparation methods. Together, these findings establish BMPDs as a quantitative framework for assessing the biochemical organization of the bone extracellular matrix.

The principal conceptual advance of this work is the shift from average biochemical measurements toward a distribution-based characterization of bone matrix composition. Our results indicate that an analogous distribution-based framework of qBEI can be applied to Raman-derived biochemical parameters describing both the mineral and organic compartments of bone. Importantly, BMPD width provides a quantitative descriptor of biochemical heterogeneity, thereby complementing conventional average measurements and capturing an additional dimension of bone material quality. In this respect, BMPDs may represent for Raman microspectroscopy what BMDDs have become for qBEI: a standardized framework that transforms thousands of local compositional measurements into biologically meaningful descriptors of tissue organization, remodeling history, and bone material quality. During the last few years, Raman microspectroscopy has progressively evolved from a research tool for measuring isolated compositional indices toward a broader technique for assessing bone material quality. Recent studies have demonstrated its ability to identify biochemical alterations associated with osteoporosis and predict bone mechanical competence using both conventional and spatially offset Raman spectroscopy (SORS) ^(21–23)^.

However, virtually all these studies rely on average spectral descriptors extracted from selected regions of interest. Our work complements these developments by introducing a framework that explicitly incorporates the spatial heterogeneity of bone tissue into Raman analysis.

An important methodological finding is that the distributions obtained for all investigated Raman parameters were predominantly unimodal and accurately modeled by gaussian functions. Although gaussian fitting was primarily adopted to maintain conceptual consistency with BMDD methodology rather than to imply an underlying biological mechanism, the excellent goodness-of-fit observed for most parameters indicates that gaussian descriptors adequately summarize the overall organization of the bone matrix. Similar to BMDD analysis, the peak, mean, and distribution width provide intuitive descriptors that can easily be compared between individuals or pathological conditions while avoiding the complexity of analyzing thousands of individual Raman spectra. A major practical advantage of the proposed workflow is the use of one-dimensional Raman line scans instead of conventional two-dimensional hyperspectral mapping. Full Raman mapping has long been recognized as the main limitation preventing routine application of Raman microspectroscopy to bone biopsies because acquisition times are prohibitively long ^(24–26)^. Here, we demonstrate that a single line scan crossing the entire trabecular thickness preserves essentially the same quantitative information as complete two-dimensional acquisitions for both organic and mineral matrix parameters, while reducing acquisition time drastically. This dramatic reduction substantially improves the feasibility of Raman-based tissue characterization for larger experimental studies and potentially for future clinical applications. The observation that a single full-thickness trabecular scan is sufficient to achieve excellent reproducibility further supports the efficiency of this simplified acquisition strategy.

Another important aspect of this study is the demonstration that BMPDs can be generated from both undecalcified pMMA-embedded bone and routinely processed decalcified paraffin-embedded specimens. Because clinical pathology laboratories almost exclusively archive decalcified paraffin blocks, the ability to derive robust biochemical descriptors from these materials considerably expands the accessibility of Raman-based analyses. Although decalcification had been linked to changes in the absolute intensity of a Raman peak ^(27)^, the absence of significant differences between rapidly decalcified pathology specimens and experimentally EDTA-decalcified pMMA sections in the present study suggests that the decalcification process itself has only a limited influence on the investigated BMPD-related Raman parameters. Consequently, BMPDs may enable retrospective studies using the extensive collections of archived paraffin bone biopsies available in pathology departments.

The small differences observed for the Hyp/Pro, CML/CH_2_, PEN/CH_2_ and GAG/Amide III ratios may not necessarily reflect true biochemical differences between decalcified paraffin-embedded and undecalcified pMMA embedded preparations. Besides potential effects of fixation, embedding and decalcification, changes in the optical properties of mineralized and demineralized bone, including refractive index mismatch, photon scattering, and Raman sampling volume, may alter the relative collection efficiency of weaker Raman bands ^(3,28–30)^. Because these parameters rely on comparatively low-intensity spectral features, they are likely to be more sensitive to such optical effects than the major collagen bands, which remained remarkably consistent across preparation methods. Consequently, the reference BMPDs established for undecalcified pMMA-embedded bone and decalcified paraffin-embedded bone should be regarded as preparation-specific and should not be used interchangeably. Rather than representing a limitation of the BMPD concept itself, these observations emphasize the need to compare pathological specimens with reference distributions obtained using identical tissue processing protocols and anatomical sampling conditions. More generally, the present study was performed using a single Raman microspectrometer under standardized acquisition conditions. Whether absolute BMPD descriptors remain invariant across different Raman platforms, including differences in optical configuration, objective numerical aperture, excitation laser characteristics, diffraction grating, detector technology, and spectral preprocessing pipelines, remains to be established. Such instrumental factors are known to influence spectral intensity, spectral resolution, and signal-to-noise ratio, potentially affecting quantitative Raman-derived parameters. Consequently, future multicenter studies will be required to evaluate the inter-instrument reproducibility of BMPDs and determine whether universal reference distributions can be established or whether instrument-specific reference datasets, analogous to those used for BMDD analysis in qBEI ^(31)^, will be required. Regardless of whether preparation- or instrument-specific calibration proves necessary, the conceptual framework of BMPDs is independent of the acquisition platform. As for BMDD analysis, only the reference distributions, not the analytical methodology itself, would require standardization for a given tissue preparation and Raman instrumentation.

The reference BMPDs generated in this study also provide the first normative database describing the distribution of multiple Raman-derived bone material properties in human bone. Interestingly, biochemical parameters reflecting collagen secondary structure, collagen maturity, proline hydroxylation, and mineral crystallinity exhibited relatively limited interindividual variability, whereas advanced glycation end-product (AGE)-related parameters and glycosaminoglycan-associated measurements displayed substantially greater biological dispersion. These observations likely reflect the distinct biological regulation of these matrix characteristics. Whereas collagen organization and mineral crystal maturation are tightly controlled during bone formation, non-enzymatic glycation and extracellular matrix remodeling are influenced by numerous metabolic and environmental factors. Furthermore, BMPD_width consistently exhibited greater biological variability than BMPD_mean or BMPD_peak, suggesting that interindividual differences in tissue heterogeneity may represent an additional dimension of bone quality that is largely overlooked by conventional average measurements.

An interesting observation was that cortical bones generally exhibited narrower BMPDs than trabecular bone despite similar average biochemical values. This finding is biologically plausible given the lower remodeling rate of cortical bone, resulting in a more homogeneous distribution of tissue ages and therefore reduced biochemical heterogeneity ^(32)^. In contrast, trabecular bone undergoes more frequent remodeling, producing a mosaic of younger and older BSUs with distinct matrix compositions. Thus, BMPD_width may provide an indirect quantitative surrogate for the spatial consequences of bone remodeling dynamics, analogous to the relationship previously established between Ca_width_ and mineralization heterogeneity.

Although the present work primarily focuses on methodological validation, the illustrative analyses of pathological bone biopsies demonstrate the potential clinical utility of BMPDs. Rather than relying on isolated mean Raman values, pathological specimens can be interpreted relative to reference distributions, allowing visualization of shifts in average composition together with alterations in tissue heterogeneity. The accompanying Z-score representation further facilitates multidimensional comparison across numerous biochemical parameters simultaneously. Such an approach could ultimately provide a biochemical fingerprint of bone tissue that complements conventional histomorphometry, qBEI, Fourier-transform infrared spectroscopy, and clinical imaging techniques. Whether specific metabolic bone diseases exhibit characteristic BMPD signatures remains an important question for future investigation.

Several limitations should nevertheless be acknowledged. First, the study was designed as a methodological validation rather than a clinical investigation, and both reference cohorts remain relatively modest in size despite encompassing broad adult age ranges. Larger multicenter studies will be required to establish definitive reference intervals and evaluate potential population-specific variability. Second, Raman parameters were calculated using peak intensity ratios, which remain indirect surrogates of biochemical composition and may be influenced by spectral preprocessing choices. Although a standardized preprocessing pipeline was implemented and technical reproducibility proved excellent, future work may benefit from multivariate spectral approaches or machine-learning-based feature extraction. Third, BMPDs were generated from linear sampling across trabeculae and cortical walls. While our validation demonstrates that this strategy adequately captures the biochemical information obtained from two-dimensional mapping, it does not fully characterize three-dimensional tissue heterogeneity. Finally, the pathological applications presented here are illustrative only and should not be interpreted as evidence of diagnostic performance.

In conclusion, we present a quantitative Raman methodology that introduces BMPDs as a new framework for characterizing the biochemical organization of bone tissue. BMPDs extend conventional Raman analyses by incorporating spatial heterogeneity into the assessment of bone matrix composition, while remaining compatible with practical acquisition times and routine histological preparations. By establishing reference distributions for multiple Raman-derived material properties and demonstrating their robustness and reproducibility, this study provides the methodological basis for future investigations exploring the role of matrix heterogeneity in bone physiology and metabolic bone disease.

## 5. Author contributions

Conceptualization: G. Mabilleau, Formal analysis: G. Mabilleau, Investigation: N. Gaborit, S. Lemiere & A. Mieczkowska, Resources: G. Mabilleau, Data curation: G. Mabilleau, Writing – original draft: G. Mabilleau, Writing – Review and editing: N. Gaborit, S. Lemiere & A. Mieczkowska, Funding acquisition: G. Mabilleau.

## 6. Acknowledgements

The authors thank the HiMolA platform – Inserm UMR_S 1229/SFR ICAT 4208 – University of Angers for support with bone assessment.

