## Supplementary materials for "Bone Material Properties Distributions from quantitative Raman analysis: a distribution-based approach to human bone matrix characterization"

Supplementary material

**Supplementary Figure S1. Workflow for the generation of BMPDs from qRA measurements in decalcified osteomedullary biopsy embedded in paraffin.** Representative Raman-derived compositional profiles were first obtained along a microscale acquisition line, providing pixel-wise measurements of key bone material parameters, including 1670/1690, 1670/1640, CML/CH2, PEN/CH2, Hyp/Pro, and GAG/Amide III ratios. Pixel-wise values were subsequently compiled into frequency distributions representing the heterogeneity of bone material properties within the analyzed region. Each distribution was modeled using gaussian fitting to extract quantitative descriptors of the BMPDs, including the coefficient of determination ($R^2$), BMPD_mean, BMPD_peak, and BMPD_width.

**Supplementary Figure S2. Workflow for the generation of BMPDs from qRA measurements in cortices of undecalcified transiliac bone biopsy embedded in pMMA.** Representative Raman-derived compositional profiles were first obtained along a microscale acquisition line, providing pixel-wise measurements of key bone material parameters, including 1670/1690, 1670/1640, CML/CH2, PEN/CH2, Hyp/Pro, GAG/Amide III, PO4/Amide I, PO4/Amide III, PO4/CH2, PO4/Hyp+Pro, CO3/PO4 ratios and mineral crystallinity. Pixel-wise values were subsequently compiled into frequency distributions representing the heterogeneity of bone material properties within the analyzed region. Each distribution was modeled using gaussian fitting to extract quantitative descriptors of the BMPDs, including the coefficient of determination ($R^2$), BMPD_mean, BMPD_peak, and BMPD_width.

**Supplementary Figure S3. Workflow for the generation of BMPDs from qRA measurements in trabecula of undecalcified transiliac bone biopsy embedded in pMMA.** Representative Raman-derived compositional profiles were first obtained along a microscale acquisition line, providing pixel-wise measurements of key bone material parameters, including 1670/1690, 1670/1640, CML/CH2, PEN/CH2, Hyp/Pro, GAG/Amide III, PO4/Amide I, PO4/Amide III, PO4/CH2, PO4/Hyp+Pro, CO3/PO4 ratios and mineral crystallinity. Pixel-wise values were subsequently compiled into frequency distributions representing the heterogeneity of bone material properties within the analyzed region. Each distribution was modeled using gaussian fitting to extract quantitative descriptors of the BMPDs, including the coefficient of determination ($R^2$), BMPD_mean, BMPD_peak, and BMPD_width.

**Supplementary Figure S4. Comparison of Raman spectral descriptors between paraffin-embedded and EDTA-treated pMMA-embedded trabecular bone.** Violin plots showing the distribution of BMPD_mean, BMPD_peak, and BMPD_width for the 1670/1690, 1670/1640, Hyp/Pro, CML/CH2, PEN/CH2, and GAG/Amide III ratios in formalin-decalcified, paraffin-embedded bone biopsies versus EDTA-decalcified, pMMA-embedded bone biopsies.

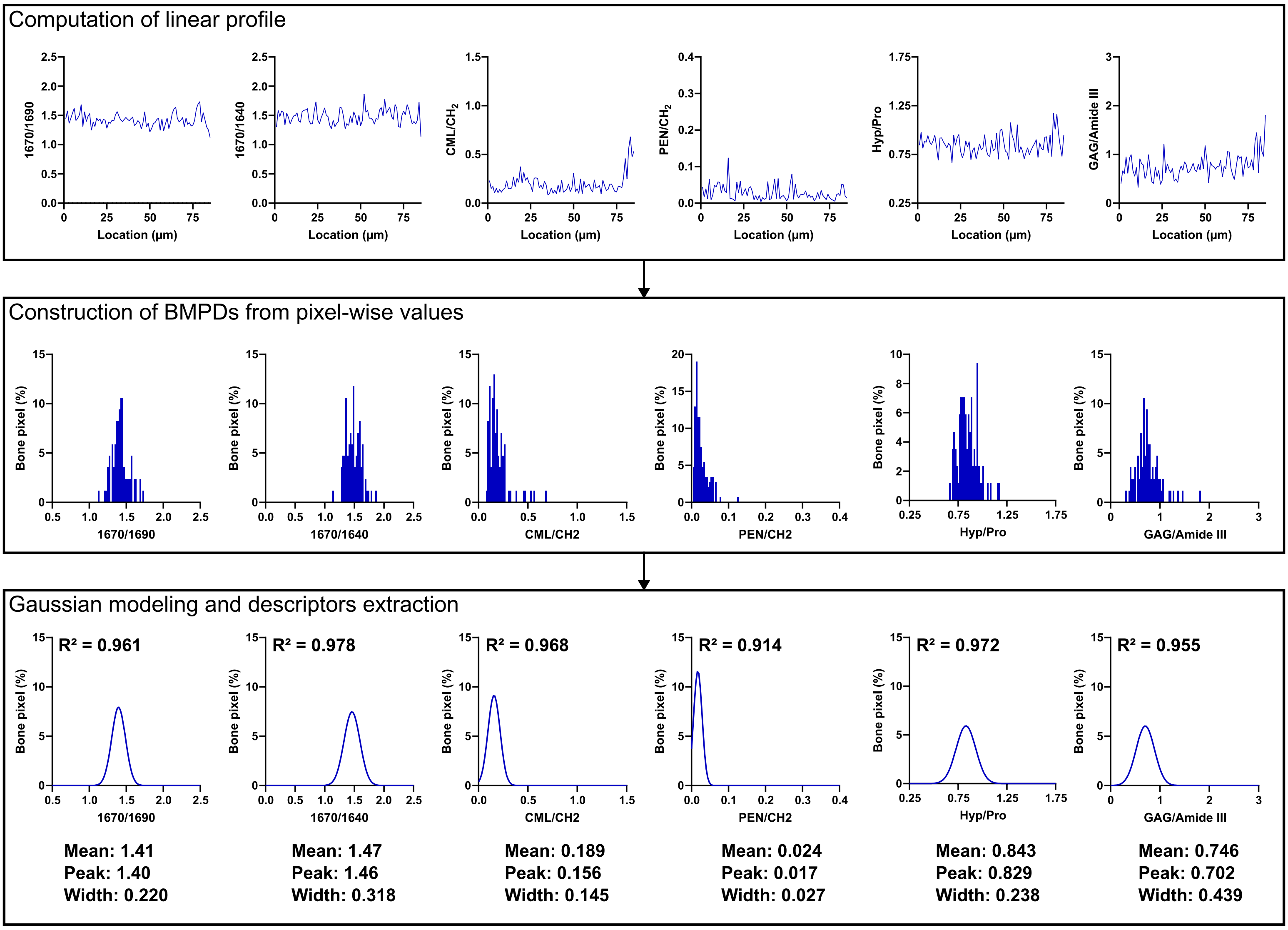


**Supplementary Figure S1**

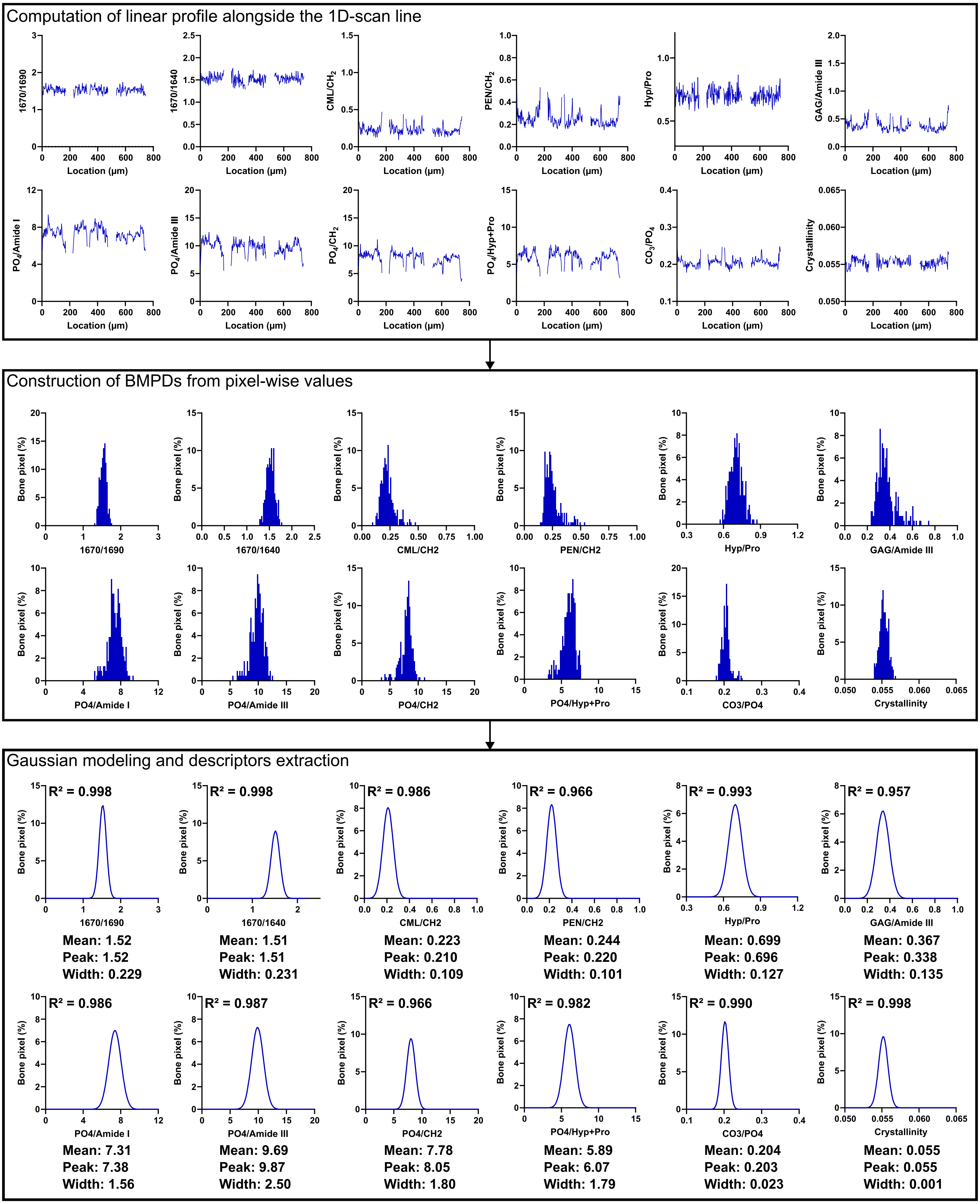


**Supplementary Figure S2**

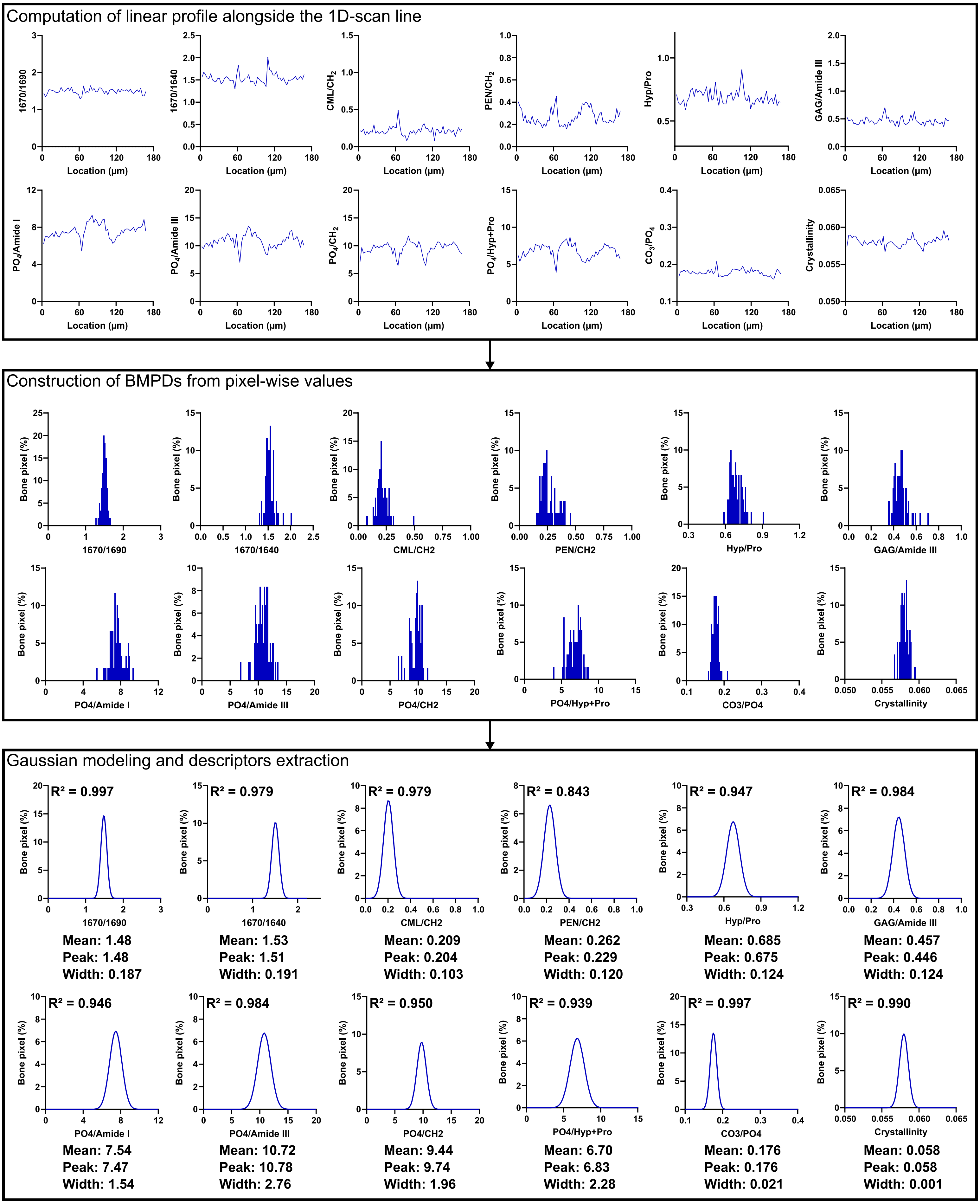


**Supplementary Figure S3**

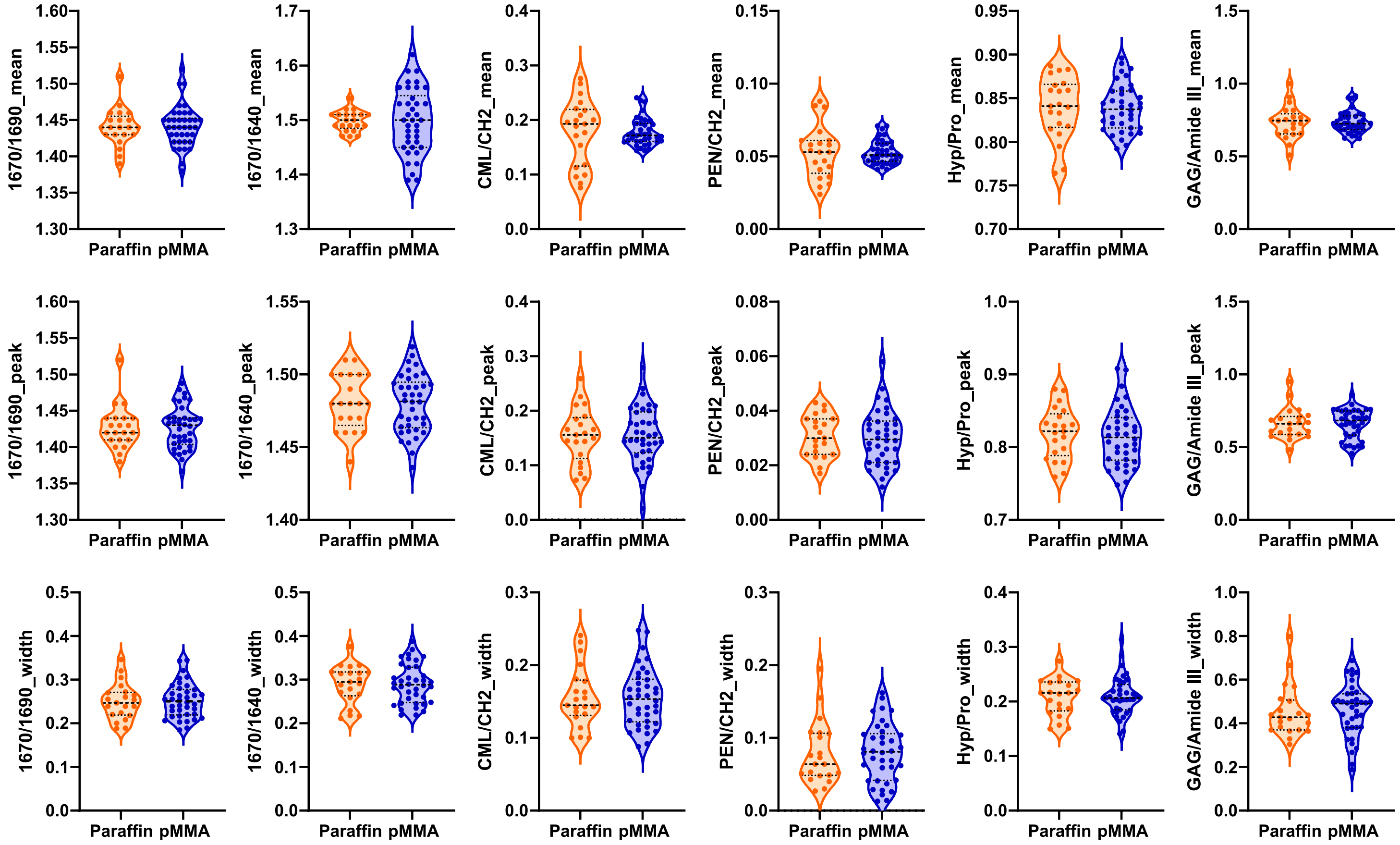
1670/1690_mean
1670/1640_mean
CML/CH2_mean
PEN/CH2_mean
Hyp/Pro_mean
GAG/Amide III_mean
1670/1690_peak
1670/1640_peak
CML/CH2_peak
PEN/CH2_peak
Hyp/Pro_peak
GAG/Amide III_peak
1670/1690_width
1670/1640_width
CML/CH2_width
PEN/CH2_width
Hyp/Pro_width
GAG/Amide III_width
Paraffin pMMA

### Figure S4

**Supplementary Table 1: Goodness of Gaussian modeling for Raman-derived BMPDs.** Coefficient of determination ($R^2$) values obtained from Gaussian fitting of BMPDs are reported for each Raman-derived bone material property according to tissue preparation and anatomical compartment. Values are presented as mean ± SD of $R^2$. Analyses are reported separately for decalcified paraffin-embedded trabecular bone, undecalcified PMMA-embedded trabecular bone, and undecalcified PMMA-embedded cortical bone. The $R^2$ values provide an assessment of the agreement between experimental BMPDs and the Gaussian model and were used to characterize the distributional behavior of each Raman-derived parameter.

| | Paraffin-embedded trabecular bone | pMMA-embedded trabecular bone | pMMA-embedded cortical bone |
|---|---|---|---|
| 1670/1690 | 0.967 ± 0.020 | 0.986 ± 0.010 | 0.994 ± 0.008 |
| 1670/1640 | 0.971 ± 0.016 | 0.961 ± 0.029 | 0.988 ± 0.013 |
| Hyp/Pro | 0.964 ± 0.019 | 0.923 ± 0.055 | 0.978 ± 0.018 |
| CML/$CH_2$ | 0.972 ± 0.027 | 0.929 ± 0.053 | 0.978 ± 0.019 |
| PEN/$CH_2$ | 0.852 ± 0.077 | 0.888 ± 0.078 | 0.956 ± 0.035 |
| GAG/Amide 3 | 0.923 ± 0.038 | 0.876 ± 0.069 | 0.953 ± 0.043 |
| v1$PO_4$/Amide I | - | 0.895 ± 0.060 | 0.948 ± 0.035 |
| v1$PO_4$/Amide III | - | 0.921 ± 0.043 | 0.967 ± 0.026 |
| v1$PO_4$/$CH_2$ | - | 0.881 ± 0.073 | 0.956 ± 0.038 |
| v1$PO_4$/Hyp+Pro | - | 0.945 ± 0.035 | 0.970 ± 0.032 |
| Crystallinity | - | 0.961 ± 0.036 | 0.990 ± 0.008 |
| v1$CO_3$/v1$PO_4$ | - | 0.957 ± 0.083 | 0.992 ± 0.006 |

**Supplementary Table 2. Agreement analysis between 2D ROI and 1D line-scan Raman acquisitions in paraffin-embedded specimen.** BMPD-derived parameters (mean, peak, and width) representing Raman-derived biochemical indices measured in decalcified paraffin-embedded bone biopsies were compared between conventional 2D ROI and reduced-dimensionality 1D line-scan acquisitions. Agreement between acquisition modalities was assessed using Bland–Altman analysis, including mean bias, standard deviation of the differences (SD), and lower and upper limits of agreement (Lower LoA and Upper LoA). Spearman's rank correlation coefficient (ρ) was additionally reported to evaluate the association between both acquisition modalities. Intraclass correlation coefficient (ICC) with its 95% confidence interval (95% CI) was calculated to assess absolute agreement between methods.

| Parameter | Mean bias | SD | Lower LoA | Upper LoA | Spearman's ρ | ICC | 95% CI ICC |
|---|---|---|---|---|---|---|---|
| **Mean** | | | | | | | |
| 1670/1690 | -0.001 | 0.007 | -0.014 | 0.013 | 0.968 | 0.948 | 0.877-0.978 |
| 1670/1640 | -0.001 | 0.009 | -0.019 | 0.016 | 0.918 | 0.871 | 0.711-0.946 |
| Hyp/Pro | -0.000 | 0.001 | -0.002 | 0.003 | 0.999 | 0.999 | 0.998-1.000 |
| CML/CH2 | 0.001 | 0.004 | -0.007 | 0.009 | 0.998 | 0.997 | 0.993-0.999 |
| PEN/CH2 | -0.000 | 0.002 | -0.004 | 0.003 | 0.995 | 0.995 | 0.987-0.998 |
| GAG/Amide III | 0.000 | 0.003 | -0.006 | 0.007 | 0.999 | 1.000 | 0.999-1.000 |
| **Peak** | | | | | | | |
| 1670/1690 | 0.002 | 0.015 | -0.027 | 0.032 | 0.740 | 0.834 | 0.638-0.929 |
| 1670/1640 | 0.006 | 0.014 | -0.022 | 0.034 | 0.696 | 0.730 | 0.443-0.881 |
| Hyp/Pro | -0.004 | 0.019 | -0.042 | 0.034 | 0.851 | 0.848 | 0.668-0.935 |
| CML/CH2 | 0.014 | 0.019 | -0.022 | 0.051 | 0.937 | 0.912 | 0.647-0.970 |
| PEN/CH2 | 0.012 | 0.014 | -0.015 | 0.038 | 0.724 | 0.385 | -0.048-0.699 |
| GAG/Amide III | 0.029 | 0.043 | -0.055 | 0.112 | 0.830 | 0.895 | 0.659-0.962 |
| **Width** | | | | | | | |
| 1670/1690 | 0.000 | 0.003 | -0.005 | 0.006 | 0.998 | 0.998 | 0.995-0.999 |
| 1670/1640 | 0.000 | 0.004 | -0.006 | 0.007 | 0.993 | 0.997 | 0.993-0.999 |
| Hyp/Pro | 0.000 | 0.003 | -0.007 | 0.007 | 0.992 | 0.995 | 0.989-0.998 |
| CML/CH2 | -0.000 | 0.005 | -0.009 | 0.009 | 0.983 | 0.994 | 0.985-0.997 |
| PEN/CH2 | 0.001 | 0.005 | -0.008 | 0.010 | 0.982 | 0.994 | 0.986-0.998 |
| GAG/Amide III | 0.001 | 0.007 | -0.013 | 0.014 | 0.993 | 0.998 | 0.996-0.999 |

**Supplementary Table 3. Agreement analysis between 2D ROI and 1D line-scan Raman acquisitions in pMMA-embedded specimen.** BMPD-derived parameters (mean, peak, and width) representing Raman-derived biochemical indices measured in 12 undecalcified pMMA-embedded bone biopsies were compared between conventional 2D ROI and reduced-dimensionality 1D line-scan acquisitions. Agreement between acquisition modalities was assessed using Bland–Altman analysis, including mean bias, standard deviation of the differences (SD), and lower and upper limits of agreement (Lower LoA and Upper LoA). Spearman's rank correlation coefficient (ρ) was additionally reported to evaluate the association between both acquisition modalities. Intraclass correlation coefficient (ICC) with its 95% confidence interval (95% CI) was calculated to assess absolute agreement between methods.

| Parameter | Mean bias | SD | Lower LoA | Upper LoA | Spearman's ρ | ICC | 95% CI ICC |
|---|---|---|---|---|---|---|---|
| **Mean** | | | | | | | |
| 1670/1690 | 0.000 | 0.008 | -0.015 | 0.016 | 0.964 | 0.957 | 0.859-0.987 |
| 1670/1640 | 0.003 | 0.010 | -0.016 | 0.021 | 0.961 | 0.958 | 0.866-0.987 |
| Hyp/Pro | 0.000 | 0.005 | -0.010 | 0.010 | 0.973 | 0.971 | 0.902-0.992 |
| CML/CH2 | 0.002 | 0.006 | -0.010 | 0.013 | 0.989 | 0.988 | 0.961-0.996 |
| PEN/CH2 | -0.000 | 0.003 | -0.006 | 0.005 | 0.998 | 0.998 | 0.993-0.999 |
| GAG/Amide III | 0.000 | 0.005 | -0.010 | 0.010 | 0.993 | 0.993 | 0.977-0.998 |
| v1$PO_4$/$CH_2$ | -0.003 | 0.011 | -0.025 | 0.020 | 0.998 | 1.000 | 0.999-1.000 |
| v1$PO_4$/Amide I | 0.002 | 0.012 | -0.022 | 0.025 | 0.997 | 1.000 | 0.999-1.000 |
| v1$PO_4$/Amide III | 0.003 | 0.011 | -0.020 | 0.025 | 0.997 | 1.000 | 0.999-1.000 |
| v1$PO_4$/Hyp+Pro | -0.002 | 0.013 | -0.028 | 0.025 | 0.998 | 1.000 | 0.998-1.000 |
| Crystallinity | -0.000 | 0.000 | -0.001 | 0.000 | 0.877 | 0.903 | 0.698-0.971 |
| $CO_3$/$PO_4$ | -0.000 | 0.004 | -0.008 | 0.007 | 0.977 | 0.924 | 0.760-0.978 |
| **Peak** | | | | | | | |
| 1670/1690 | -0.005 | 0.016 | -0.036 | 0.025 | 0.595 | 0.719 | 0.303-0.909 |
| 1670/1640 | -0.007 | 0.026 | -0.058 | 0.045 | 0.897 | 0.938 | 0.807-0.982 |
| Hyp/Pro | -0.004 | 0.009 | -0.021 | 0.014 | 0.806 | 0.963 | 0.879-0.989 |
| CML/CH2 | 0.001 | 0.004 | -0.008 | 0.009 | 0.997 | 0.991 | 0.970-0.997 |
| PEN/CH2 | 0.000 | 0.012 | -0.022 | 0.023 | 0.797 | 0.961 | 0.869-0.988 |
| GAG/Amide III | -0.001 | 0.007 | -0.015 | 0.014 | 0.956 | 0.983 | 0.941-0.995 |
| v1$PO_4$/$CH_2$ | -0.038 | 0.104 | -0.242 | 0.166 | 0.997 | 0.998 | 0.994-0.999 |
| v1$PO_4$/Amide I | -0.019 | 0.080 | -0.175 | 0.137 | 0.986 | 0.993 | 0.976-0.998 |
| v1$PO_4$/Amide III | -0.002 | 0.130 | -0.256 | 0.253 | 0.965 | 0.989 | 0.961-0.997 |
| v1$PO_4$/Hyp+Pro | 0.015 | 0.075 | -0.132 | 0.162 | 0.986 | 0.993 | 0.978-0.998 |

| | | | | | | | |
|---|---|---|---|---|---|---|---|
| Crystallinity | -0.000 | 0.001 | -0.001 | 0.001 | 0.514 | 0.585 | 0.086-0.857 |
| $CO_3/PO_4$ | 0.001 | 0.006 | -0.011 | 0.012 | 0.765 | 0.836 | 0.524-0.950 |
| **Width** | | | | | | | |
| 1670/1690 | 0.000 | 0.002 | -0.004 | 0.004 | 0.998 | 0.999 | 0.998-1.000 |
| 1670/1640 | 0.000 | 0.002 | -0.003 | 0.004 | 0.998 | 1.000 | 0.999-1.000 |
| Hyp/Pro | -0.000 | 0.003 | -0.005 | 0.005 | 0.991 | 0.999 | 0.998-1.000 |
| CML/CH2 | -0.001 | 0.003 | -0.007 | 0.006 | 0.993 | 0.999 | 0.996-1.000 |
| PEN/CH2 | -0.002 | 0.007 | -0.016 | 0.012 | 0.993 | 0.994 | 0.979-0.998 |
| GAG/Amide III | 0.000 | 0.003 | -0.006 | 0.006 | 0.993 | 1.000 | 0.999-1.000 |
| $v1PO_4/CH_2$ | 0.000 | 0.017 | -0.033 | 0.033 | 0.998 | 1.000 | 0.999-1.000 |
| $v1PO_4$/Amide I | -0.005 | 0.014 | -0.032 | 0.022 | 0.998 | 1.000 | 0.999-1.000 |
| $v1PO_4$/Amide III | 0.002 | 0.016 | -0.029 | 0.033 | 0.992 | 1.000 | 0.999-1.000 |
| $v1PO_4$/Hyp+Pro | -0.003 | 0.022 | -0.045 | 0.040 | 0.998 | 0.999 | 0.998-1.000 |
| Crystallinity | 0.000 | 0.000 | -0.000 | 0.000 | 0.914 | 0.946 | 0.824-0.984 |
| $CO_3/PO_4$ | 0.000 | 0.000 | -0.001 | 0.001 | 0.986 | 0.997 | 0.990-0.999 |
